# Directional Tuning of Phase Precession Properties in the Hippocampus

**DOI:** 10.1101/2021.12.08.471702

**Authors:** Yuk-Hoi Yiu, Jill K. Leutgeb, Christian Leibold

**Author notes:** **Corresponding author:** Christian Leibold, Albert-Ludwigs-Universität Freiburg, Bernstein Center Freiburg, Hansastr. 9a, D-79104 Freiburg, Germany.

## Abstract

Running direction in the hippocampus is encoded by rate modulations of place field activity but also by spike timing correlations known as theta sequences. Whether directional rate codes and the directionality of place field correlations are related, however, has so far not been explored and therefore the nature of how directional information is encoded in the cornu ammonis remains unresolved. Here, using a previously published dataset that contains the spike activity of rat hippocampal place cells in the CA1, CA2 and CA3 subregions during free foraging of male Long-Evans rats in a 2D environment, we found that rate and spike timing codes are related. Opposite to a place field’s preferred firing rate direction spikes are more likely to undergo theta phase precession and, hence, more strongly impact paired correlations. Furthermore, we identified a subset of field pairs whose theta correlations are intrinsic in that they maintain the same firing order when the running direction is reversed. Both effects are associated with differences in theta phase distributions, and are more prominent in CA3 than CA1. We thus hypothesize that intrinsic spiking is most prominent when the directionally modulated sensory-motor drive of hippocampal firing rates is minimal, suggesting that extrinsic and intrinsic sequences contribute to phase precession as two distinct mechanisms.

## 1 Introduction

Hippocampal place cells establish a neuronal representation of space by exhibiting elevated firing rates at only few locations in an environment, called place fields (O’Keefe and Dostrovsky, 1971). Place field firing is thought to underlie an animal’s capacity to navigate in space and to form spatial memories (Morris et al., 1982; Moser et al., 1993; Nakazawa et al., 2002). Place field firing also includes a temporal code associated with the theta oscillation (4-12 Hz) of the local field potential (O’Keefe and Recce, 1993): As a rat passes through a place field, the spikes phase-precess, i.e., they occur at successively earlier theta phases thereby encoding the relative location of the animal within a place field (O’Keefe and Recce, 1993; Harris et al., 2002). Phase precession is generally thought to implement a compression of behavioral sequences to the theta time scale, such that, in a 100 ms time window, spikes of multiple place cells are elicited in the same order as the activation of the associated place fields along the animal’s trajectory over the time scale of seconds (Melamed et al., 2004; Dragoi and Buzsaki, 2006; Foster and Wilson, 2007; Jaramillo and Kempter, 2017).

In one-dimensional mazes, the trajectory of an animal can be uniquely mapped to a sequence of place fields, thus, spike sequences on the theta time scale (Foster and Wilson, 2007) can be easily interpreted as reflecting memories of previous locations or planning of future actions (Feng et al., 2015). In two-dimensional environments, place fields can be entered from multiple directions and, hence, place cells generally take part in encoding multiple trajectories. Thus place cells could either be linked to multiple sequences or place field sequences could be directional. In the former case sequential structure would be imposed by sensory-motor (extrinsic) inputs, whereas the latter case would render sequences of intrinsic origin supported by recurrent circuits or associative loops. Previous reports revealed that pair correlation lags of place fields in the CA1 subregion depend on running direction (Huxter et al., 2008) and thus support the extrinsic hypothesis, but similar analyses for the CA2 and CA3 subregions that differ substantially in their cytoarchitecture, plasticity, and protein chemistry are missing.

In addition to sequence order, directional information is also available to the entorhinal-hippocampal circuits from head direction cells of the postsubiculum (Taube et al., 1990a; Taube et al., 1990b) and the medial entorhinal cortex (Sargolini et al., 2006; Giocomo et al., 2014), and to a limited extent from within the cornu ammonis itself (Leutgeb et al., 2000). These inputs might explain observed directionality of some place fields (Leutgeb et al., 2004; Acharya et al., 2016; Mankin et al., 2019), however, it is unclear to which extent this rate directionality is related to directionality of theta sequences.

Past studies have shown that, compared to the CA1 region, place cells in CA3 demonstrate a more persistent and consistent activity pattern over an extended period of time (Mankin et al., 2012) and across cue-altered environments (Lee et al., 2004), as well as a higher stability of place field dynamics across multiple recording sessions in the same environment (Mizuseki et al., 2012). Consistently, CA3 place representations stabilized more slowly after a change of environment than in CA1 (Leutgeb et al., 2004). The stability, consistency and slower stabilization of CA3 place fields, in addition to the classical anatomy indicating strong recurrent connectivity (Amaral and Witter, 1989; Ishizuka et al., 1990), lead to a belief that CA3 activity patterns are more reliant on internal network dynamics and less influenced by external sensory inputs. Therefore, we hypothesized that the theta sequence activity in CA3 should be less dependent on the directionality in the animal’s behaviours than CA1.

Our results show that, although the extrinsic contribution to theta scale firing is dominant in all subareas, this is indeed least visible in CA3. Moreover, we observe directionality in the phase precession properties such that CA3 displays later spike phases in the running direction opposite to the best firing rate direction.

## 2 Materials and Methods

### Experimental Design and Statistical Analysis

We re-analyzed a previously published data set in Mankin et al. (2012); Mankin et al. (2015). For detailed description of the data collection we refer to the original papers. In brief, the data set involves eight male Long-Evans rats who were trained to forage for randomly scattered cereal crumbs in either a square (80cm × 80cm) or a 16-sided polygon (50cm radius, also referred to as circular) enclosure. The experiment began after animals were trained 9-20 days in the enclosure. Single units were recorded simultaneously from the CA1, CA2, and CA3 subregions for the course of the experiment, which lasted for two days. On each day the rats completed two blocks of four 10-minute sessions, with two sessions in the square enclosure and two sessions in the circular enclosure assigned in random order. The enclosures contain a 20cm-wide white cue card on an inside wall, and the cue card maintained a constant angle with the cues outside the room.

Data analysis and statistical tests were performed on Python using SciPy (Virtanen et al., 2020), NumPy (Harris et al., 2020) packages and custom routines based on CircStat toolbox (Philipp, 2009). We used nonparametric Kruskal-Wallis tests with post hoc Dunn’s test for statistical comparisons. Normally distributed data was tested using Student’s t-test. The Watson-Williams test was used for circular data. For categorical data, we used chi-square and Fisher’s exact tests. P-values were adjusted for multiple comparisons using Benjamini–Hochberg procedure (Benjamini and Hochberg, 1995). We used two-tailed tests throughout the paper except for binomial tests. *p* = 0.05 is chosen as the significance level.

### Place field detection and delimination

For each place cell, we computed a spatial map of firing rates from each 10-minute session by dividing the spikes counts by the occupancy time in each space bin (1cm × 1cm). Spikes and occupancy were smoothed by a Gaussian filter with a standard deviation of 3cm. Place fields were segmented along the closed contour line located at 20% of the maximum rate and accepted if the peak rate exceeds 1Hz, the field area is larger than 25cm^2^, and the average firing rate within the field is larger than outside the field. In the analysis, we separated place fields into a border and a non-border group. The groups are distinguished by the 20% line touching the boundary of the enclosure. Each place cell could have multiple place fields that were analyzed separately.

Two place fields are said to be a pair if both field areas intersect and contain at least 16 spikes of one field that occur next to a spike of the other field within a time window of ±0.08s (for illustration, see Figure 4B). We defined pairs as border pairs if at least one of the place fields touches the border. In non-border pairs, both place fields do not touch the border.

### Directionality of place fields

To obtain the directional tuning curve of place fields, we employed a Maximum Likelihood Maximization (MLM) model (Cacucci et al., 2004). In brief, the model assumes an independent relation between positional and directional firing probability distributions, whose product is the firing probability. The product assumption furthermore ensures that directionality tuning is restricted to the place field and that the tuning depends on a weighted sum of position and direction. The solution of the directional term, which is also the estimated directional tuning curve, can be fit by iteratively maximizing firing likelihood to the observation of spikes. This MLM model has an advantage of reducing sampling bias, which usually arises at the enclosure borders where certain heading directions are severely under-sampled.

Significance of directionality is determined by comparing the mean resultant vector length (*R*) of the directional tuning curve to a shuffling distribution, which is obtained by randomly shifting the spike times in a cyclic fashion along a trajectory concatenated from all path segments traversing the place field. The random time shift was repeated 200 times for each place field. A place field is classified as significantly directional if the *R* value exceeds the 95-percentile of its shuffling distribution. For field pairs, the same method is used to determine the significance of directional selectivity, except that the *R* is calculated from spike pairs.

Significance of the preferred precession direction is also determined by comparison of *R* to a distribution obtained from shuffling spike times over all passes through the field. In this case the *R* value for precession directionality is computed from the directions of field traversals which exhibit phase precession. A pass is labeled as precessing if the phase-position relation of the spikes has a negative slope derived from linear-circular regression (Kempter et al., 2012).

### Correlation lags

We produced cross-correlograms for the spike trains of every field pair, with the resolution of 5 ms and a time window between -150 ms and 150 ms. The resultant cross-correlograms were then band-pass filtered (5–12 Hz) to derive the correlation lag as the phase at 0 time lag from the Hilbert transform of the theta filtered correlograms.

### Classification of extrinsic and intrinsic pairs

In order to quantify the dependence of correlation lag on directionality, we devised the measures of ‘extrinsicity’ and ‘intrinsicity’ for each field pair. The extrinsicity is computed as the Pearson’s correlation coefficient between the cross-correlogram for runs from one field to another and the cross-correlogram for runs in the opposite direction but with the sign of time-axis being flipped. The Pearson’s correlation coefficient (*r*) is then linearly transformed 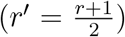 to be in the range of 0 and 1. The extrinsicity is close to 1 if a field pair was mainly driven by the external sensory input, as the sign of correlation lag would be reversed if the place fields were traversed in a reversed order. Similarly, the intrinsicity is computed as the Pearson’s correlation coefficient between the two cross-correlograms without flipping the sign of time-axis. The value of intrinsicity is close to 1 if the correlation lag of a field pair was mainly dependent on its intrinsic dynamics but less on the external sensory-locomotor input, leading to similar correlograms in two running directions. Note that, using this definition, extrinsicity and intrinsicity are two independent values that are not necessarily correlated. We classify a field pair as “extrinsic” if its extrinsicity exceeds its intrinsicity, and as “intrinsic” if its intrinsicity exceeds its extrinsicity.

### Inclusion criteria for analysis

The animal trajectory is split to passes entering and exiting the place fields or pairs. Intervals in which the speed of animal is below 5cm/s were excluded. The passes are then chunked to smaller segments in which the speed was always above 5cm/s.

The pass segments and their spikes are only included if the pass duration is longer than 0.4 s and satisfies a straightness threshold, i.e.,

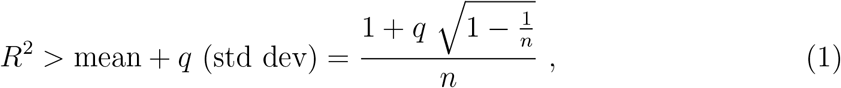

where *q* = 5, *R* is Rayleigh Vector length of the animal’s heading samples, and *n* is number of heading samples. In case the pass is chunked due to low speed, in order to determine the travelled distance of the animal inside the place field relative to the entry point, only the first pass segment entering the place field is included for analysis. The relative position of the animal can thus be determined by distance travelled divided by the field diameter.

Pair-crossing passes are classified as *A* → *B* if they start from an area within field A but not field B, and end in an area within field B but not field A. The opposite criteria apply for *B* → *A* passes. Passes that do not satisfy the above criteria, or cross either one of the field boundaries more than once, are not assigned to any of the directional groups (*A* → *B* or *B* → *A*). These unassigned passes were also included in computing the firing rate directionality, but excluded for cross-correlation analyses. As a result, field pairs that have no spike-pairs along the *A* → *B* or *B* → *A* passes were further excluded in the cross-correlation analyses.

### Model simulation

Romani and Tsodyks (2015) proposed a recurrent network model of the hippocampus, which we adapted and simulated using our trajectory data. Here we briefly summarize the key equations of the model. The dynamics of firing rate *m*_*i*_(*t*) of place cell *i* at time *t* is given by

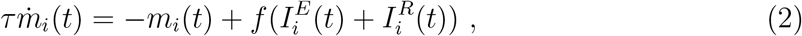

where *τ* = 10ms is the time constant, *f* (.) is the firing rate function, 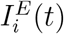 is the sum of external positional and theta oscillatory inputs and 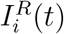 is the recurrent input from the other neurons. The latter is computed as

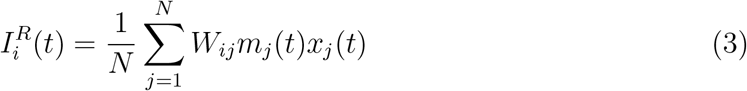

with *W*_*ij*_ denoting the synaptic weight from neuron *j* to *i* and *x*_*j*_ denoting the depletion state of the synaptic vesicle pool, which is recovering with time constant *τ*_*R*_ = 800ms, i.e.,

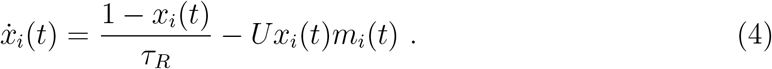

The parameter 0 *< U* ≤ 1 denotes the release probability. The introduction of *x*_*i*_(*t*) penalizes the recurrent input from the highly activated place cells with a delay, and therefore, it produces asymmetrical weight couplings which are stronger in the forward direction as the animal moves.

We adopted the model parameters from the toroidal environment described in Romani and Tsodyks (2015). The periodicity of the environment was removed by clipping the cosine function cos(*x*) at the value of -1 for |*x*| *> π*. Our simulation thus has 32 × 32 neurons equally spaced across a 2*π* × 2*π* unit squared environment. We randomly chose one recording session in the square arena from the experimental data as the trajectory, and re-scaled the trajectory into the range of 0 and 2*π* and the average speed to be the same as in the original study (2*π/*5 unit per second). Simulation was implemented with 1 ms temporal resolution using the Euler method. Spikes from each place cell were then sub-sampled by a fraction such that the average spike count of all simulated place fields is the same as the experimental data.

## 3 Results

### 3.1 Directional tuning of place cells

Directionality of hippocampal place field activity has been reported in a number of previous studies (Leutgeb et al., 2004; Acharya et al., 2016; Mankin et al., 2019) but quantitative comparisons between those studies were hampered because of the use of different methods and behavioral paradigms. Here we analyze the firing properties of simultaneously recorded CA1, CA2, and CA3 neural networks under identical experimental conditions. We thus first applied one established rate-based directionality analysis on the data sets used in this paper (Mankin et al., 2012; Mankin et al., 2015) for further comparison. Directional tuning for each place field was quantified using the mean resultant vector length (*R*) obtained from directionality fields derived by the Maximum Likelihood Model (MLM) proposed in (Cacucci et al., 2004) (see Figure 1A for single examples and 1 B for populations). Since mean resultant vector lengths are strongly biased by the number of observations (Figure 1 B), we decided to analyse directional tuning as a function of the spike count threshold criterion for including place fields (Figure 1 C), and include only the fields with spike counts higher than 40 in our statistical analysis of place field directionality. We found that all CA regions contain a fraction of directional fields that is above chance level (Binomial test: CA1, 146/800=0.1825, *p* = 1.8*e* − 41; CA2, 111/521=0.2131, *p* = 2.4*e* − 38; CA3, 45/396=0.1136, *p* = 3.5*e* − 07). The amount of directionality in all regions not only depends on the overall spike count threshold of the place field (showing an initial increase that is expected from the gain in statistical power), but also on whether the place field is located at the boundary of the arena (Figure 1C).

**Figure 1:**
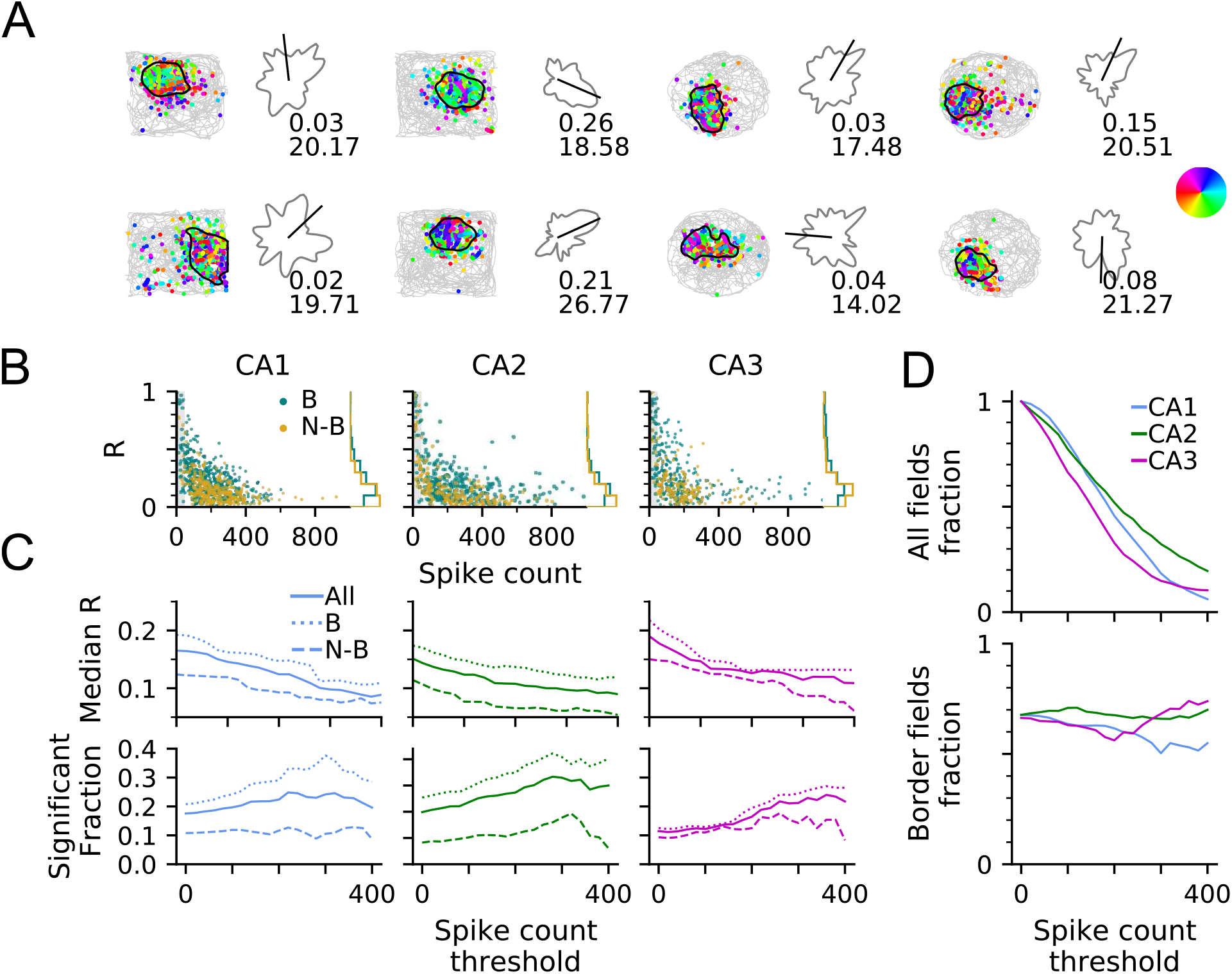
Directionality of place field firing rates. (A) Examples of place fields in square and circular enclosures of a free-foraging experiment overlaid with the trajectory of the animal in one recording session and spike events color-coded by heading direction (see color wheel at the right). Directional tuning curve is shown to the right of each spike-position plot. Mean direction (black bar), mean resultant vector length *R* (upper text), peak rate in Hz (lower text). (B) *R* and total within-field spike counts of all border (“B”, teal) and non-border (“N-B”, golden) place fields, as well as marginal distribution of *R* (right). *R* values are strongly biased by sampling. Therefore, we excluded the fields with spike counts below 40 in our directionality analysis, indicated by the shaded region. (C) Median *R* (top row) and fraction of significantly directionally tuned place fields (bottom) by different spike number thresholds for all (solid line), border (dotted line) and non-border (dashed line) fields in each brain region as indicated. CA1 and CA2 directionality is strongly border driven. (D) Fraction of all place fields (top) and border fields (bottom) by spike count thresholds.

Comparing the place field directionalities between CA regions, CA1 and CA3 have a higher median *R* than CA2 (Kruskal-Wallis test: CA1 (*n* = 800) vs CA2 (*n* = 521) vs CA3 (*n* = 396), *H*_(2)_ = 22.18, *p* = 1.5*e* − 05; *post hoc* Dunn’s test with Benjamini-Hochberg correction: CA1 vs CA2, *p* = 0.0003, CA2 vs CA3, *p* = 3.8*e* − 05, CA1 vs CA3, *p* = 0.1920). The fraction of significantly directional fields in CA3 is lower than in CA1 and CA2 (Fisher’s exact test for independence of significant fractions with Benjamini-Hochberg correction: CA1 vs CA2, *p* = 0.2656; CA1 vs CA3, *p* = 2.2*e* − 05; CA2 vs CA3, *p* = 0.0074). However, as we increase the spike count threshold to admit only fields that are highly sampled, the fraction of significantly directionally tuned fields becomes similar in CA1 and CA3, as far as our data allows such a comparison owing to the only very few CA1 and CA3 fields with high spike numbers (Figure 1D).

In order to accurately interpret the above results, we looked into possible confounds. A major influence on directionality could arise from the presence of arena boundaries, because both, they act as salient sensory landmarks and they introduce a behavioral bias. We thus further separated place fields into border and non-border fields. Including all the place fields regardless of spike counts, we found that border fields in CA1 generally have higher directional selectivity than in the non-border case (Kruskal-Wallis test: border (*n* = 537) vs nonborder (*n* = 263), *H*_(1)_ = 62.33, *p* = 2.9*e* − 15; Fisher’s exact test: *p* = 5.7*e* − 05), while CA3 exhibits significant border difference in median *R* (border (*n* = 257) vs nonborder (*n* = 139), *H*_(1)_ = 15.31, *p* = 9.1*e* − 05) but not in significant fraction (Fisher’s exact test: *p* = 0.0885). Similar to CA1, directionality measures in CA2 also exhibit a significant border effect (Kruskal-Wallis test: border (*n* = 359) vs nonborder (*n* = 162), *H*_(1)_ = 31.95, *p* = 1.6*e* − 08; Fisher’s exact, *p* = 3.6*e* − 07).

We thus conclude that place field rates in all CA areas encode running direction, and directionality in CA1 and CA2 is more strongly induced by borders, whereas this is not the case for CA3, in which the directionality is more similar between border and non-border fields. Assuming boundaries to induce a strong sensory-motor constraint, this is already a first hint that CA1 activity is more strongly influenced by extrinsic factors than CA3.

### 3.2 Preferred direction for phase precession in place fields

Since, in addition to the firing rate code, place field activity is also temporally organized on the theta scale by phase precession (Figure 2A), we also asked to what extent directionality is also reflected in this temporal code. For each place field, we therefore identified a direction in which phase precession is more likely to occur using single pass phase precession analysis (Schmidt et al., 2009; Kempter et al., 2012) (see Materials and Methods for inclusion criteria). In brief, we fit a linear-circular regression line for the phase-position relation in every single pass. Passes with negative regression slope between −2*π* and 0 (per pass length) are classified as phase precessing (Figure 2B).

**Figure 2:**
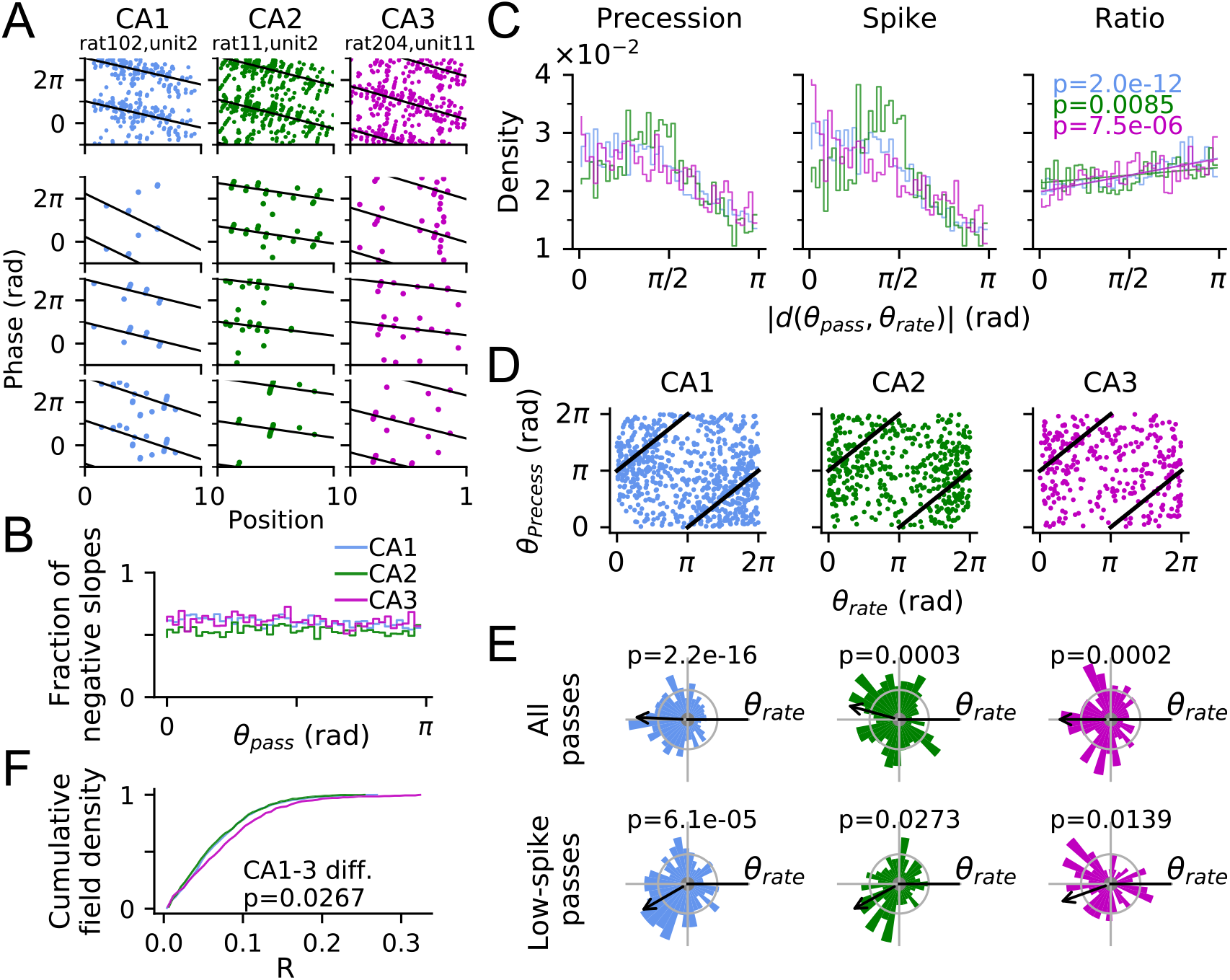
Phase precession per spike is most prevalent opposite to the direction of highest firing rate. (A) Top row: Three examples of phase precession pooled over all passes in one recording session. Position is normalized to be between 0 and 1 with respect to the moments of entering and exiting the place field. Linear-circular regression line is indicated in black, characterizing phase precession by its slope and onset phase. Bottom rows show phase precession in example passes. Passes with slope between −2*π* and 0 are classified as incidents of phase precession. (B) Fraction of fitted negative slopes among all passes as a function of pass direction *θ*_pass_ for CA1, CA2 and CA3 (color as indicated) indicates a similar amount of precessing passes in all subregions. (C) Distribution of phase precessing passes in the whole data sets (left) mirrors the elevated spike count along the fields’ preferred rate direction *θ*_rate_(middle) pooled over all precessing passes as a function of absolute angular deviation |*d*(*θ*_pass_, *θ*_rate_)| from pass direction *θ*_pass_. We therefore normalized the spike distribution by precession occurrences (ratio of left and middle graph) to obtain a distribution of phase precession per spike (right), which increases with angular distance |*d*(*θ*_pass_, *θ*_rate_)| (p-values from Spearman’s correlations: CA1, *r*_*s*_ = 0.83, CA2, *r*_*s*_ = 0.39, CA3, *r*_*s*_ = 0.62) indicating an excess of precession at *d* = *π* that cannot just be explained by increased firing. (D) The same analysis on a field-wise level shows preferred precession directions (per spike) *θ*_precess_ of single fields and best firing rate direction *θ*_rate_ to be offset by about *π* (marked by black line). (E) Top: Same as D shown as normalized polar histograms of *θ*_precess_ relative to *θ*_rate_. Bottom: only low-spike passes (25-percentile) are admitted (p-values are derived from V-test versus the Null hypothesis of a circular mean at *π*) to control for high rate bias. Arrow marks direction of the mean resultant vector of the distribution, with the best rate direction pointing to the right. (F) Cumulative distribution of *R* of all place fields in CA1, CA2 and CA3. Kruskal-Wallis test indicates strongest directionality in CA3.

First, we computed the density of phase precession occurrences from all passes in all fields, as a function of pass direction relative to the respective field’s preferred firing rate direction (Figure 2C). We find most precessing passes along the best rate direction, reflecting the fact that more spikes should give rise to more detectable phase precession. However, not all spikes may contribute to phase precession to the same degree, either because of different levels of phase noise or because they occur outside theta sequences. If phase precession is directional beyond a simple spike count effect, it needs to show in an analysis per spike. Thus computing the density of precessing passes per spike (Figure 2C right), we found that phase precession is more likely to occur the more the pass direction differs from the field’s preferred rate direction (Spearman’s correlation: CA1, *r*_*s*(6719)_ = 0.83, *p* = 2.0*e* − 12; CA2, *r*_*s*(4837)_ = 0.39, *p* = 0.0085; CA3, *r*_*s*(2625)_ = 0.62, *p* = 7.5*e* − 06), indicating that spike rate and phase precession differentially contribute to rate directionality.

In addition to this population-wide analysis, we also identified the direction in which precession is most probable per spike for each field separately and call it the best precession angle *θ*_precess_. Histograms of preferred precession angles from all place fields separately (Figure 2E top row) also demonstrate a significant *π* shift from their preferred firing direction (V-test versus *π* direction: CA1, *V*_(731)_ = 155.54, *p* = 2.2*e* − 16; CA2, *V*_(466)_ = 51.75, *p* = 0.0003; CA3, *V*_(341)_ = 47.12, *p* = 0.0002). However, by comparing the *R* values of precessing passes to a shuffling distribution (see Materials and Methods for shuffling procedures), we found that only 24/829 (2.9%), 11/560 (2.0%) and 20/441 (4.5%) of place fields in CA1, CA2 and CA3 respectively, exhibit significant preferred precession direction, which is not significant under binomial tests (CA1, *p* = 0.9990; CA2, *p* = 0.9999; CA3, *p* = 0.7034). We thus conclude that, while on the level of the single field the anti-phase relation between spike count and phase precession is weak and does not reach significance (and therefore has likely not been identified previously), there is strong indication of such a relation on the population level.

To further rule out that the *π* shift between best rate and best precession direction might arise as an epiphenomenon of different spike counts, with the opposite of the preferred rate direction being over-represented by the normalization process, we re-computed the histograms of preferred precession angles by limiting the passes to only those with low spike counts (*<* 25% quantile of all precessing passes in each CA region) such that there is no firing rate directionality left in the used data. Our results show that the place fields in all CA regions still demonstrate a significant *π* shift from the preferred rate direction (Figure 2E bottom row, V-test versus *π* direction: CA1, *V*_(294)_ = 46.58, *p* = 6.1*e* − 05; CA2, *V*_(226)_ = 20.44, *p* = 0.0273; CA3, *V*_(120)_ = 17.04, *p* = 0.0139). Thus, on the population level, the direction of best phase precession displays a consistent and significant bias towards the opposite of the direction of best firing rate corroborating the hypothesis of distinct coding schemes, and, hence, input streams, for spike timing and rate (Huxter et al., 2003).

Finally, we also compared distributions of Rayleigh vector lengths *R* for phase precession tuning and found that, whereas CA1 & CA2 seem to have similar directionality, CA3 exhibits a significantly higher directional selectivity (Figure 2F, Kruskal-Wallis test: CA1 (*n* = 753) vs CA2 (*n* = 485) vs CA3 (*n* = 363), *H*_(2)_ = 9.37, *p* = 0.0092; *post hoc* Dunn’s test with Benjamini-Hochberg correction: CA1 vs CA2, *p* = 0.3351, CA2 vs CA3, *p* = 0.0084, CA1 vs CA3, *p* = 0.0267) further suggesting that directional information in CA3 place field activity is distinct from CA1 & CA2.

### 3.3 Phase precession properties show dependence on pass direction

To further support the existence of directional effects on phase precession and to elucidate the underlying processes, we searched for directional modulations of phase precession properties by quantifying single pass onset phase and precession slope and plotted their occurrence density (Figure 3A) for different relative pass directions |*d*| (defined as the absolute circular difference between pass direction and preferred rate direction of the place field). While there was no significant correlation between precession slope and |*d*| in any CA region (CA1, *r*_(5020)_ = −0.008, *p* = 0.5768; CA2, *r*_(3514)_ = 0.031, *p* = 0.0564; CA3, *r*_(2044)_ = 0.013, *p* = 0.5558), we found that phase onsets slightly but significantly increase as the heading deviates more from the preferred rate direction in CA3 (*r*_(2625)_ = 0.078, *p* = 5.4*e* − 05), also, but barely significantly, in CA1 (*r*_(6719)_ = 0.024, *p* = 0.0459), but not significantly in CA2 (*r*_(4837)_ = −0.023, *p* = 0.0952). Thus the opposite directions of best firing rate are signified by later phases, which could reflect more prospective parts of theta sequence activity (Foster and Wilson, 2007).

**Figure 3:**
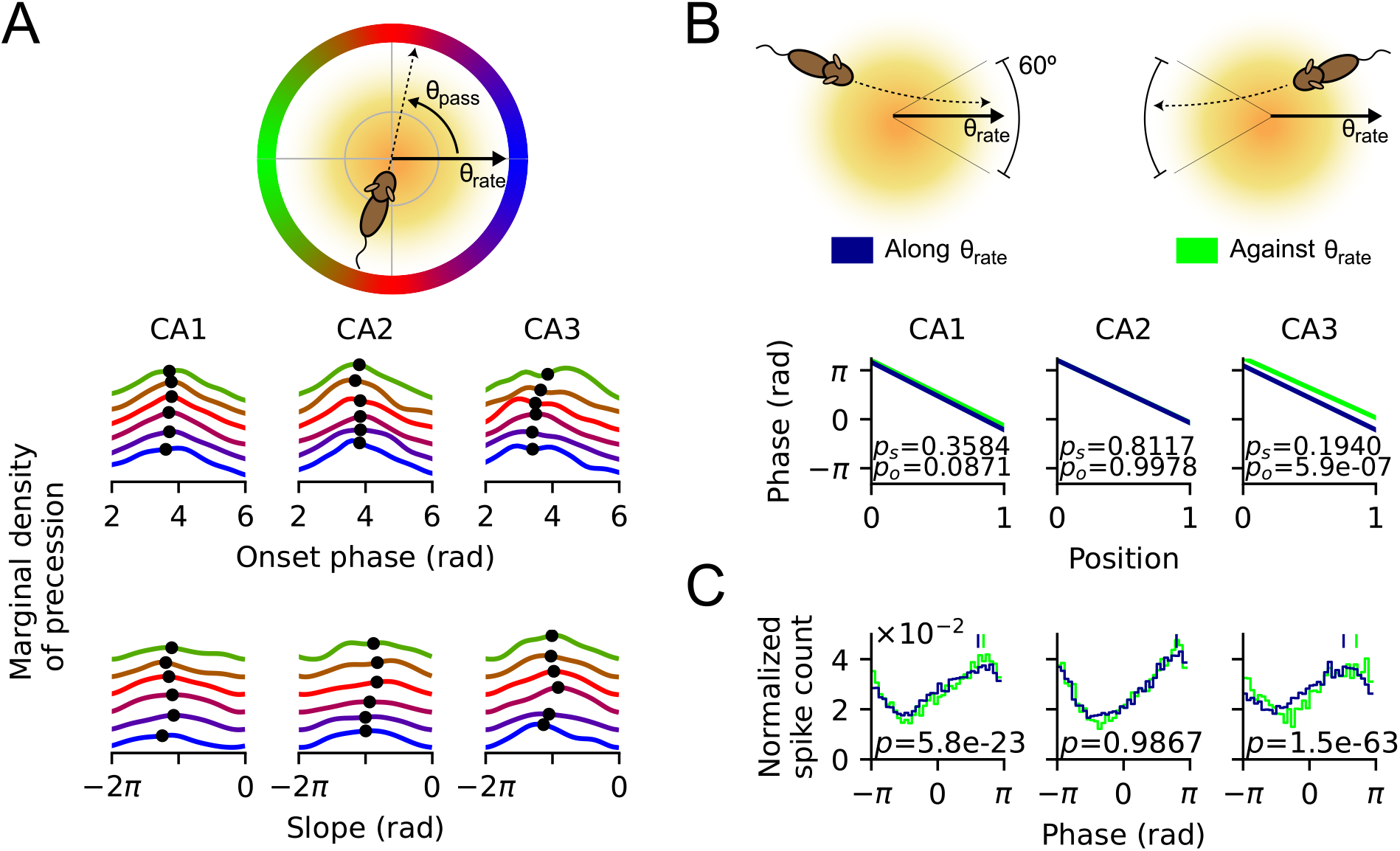
Directional dependence of phase precession properties. (A) Marginal distributions of precession density as a function of onset phase and slope, color-coded by the difference between pass angle *θ*_pass_ and best rate angle *θ*_rate_ (see illustration of the color code on top). Black dot indicates the circular mean of marginal density. (B) Average phase-position relations for cases where the animal is running along (green) and against (blue) *θ*_rate_. See schematic illustration above. Against-*θ*_rate_ condition has higher onset than along-*θ*_rate_ condition. *p*_*o*_ denotes the p-value from Watson-Williams test for onset difference. [along-*θ*_rate_ vs against-*θ*_rate_, mean ± SEM in radians: CA1, 3.66 ± 0.04 vs 3.77 ± 0.05, *F*_(1,2073)_ = 2.93, *p* = 0.0871; CA2, 3.78 ± 0.04 vs 3.78 ± 0.05, *F*_(1,1301)_ = 0.00, *p* = 0.9978; CA3, 3.44 ± 0.06 vs 3.94 ± 0.07, *F*_(1,848)_ = 25.34, *p* = 5.9*e* − 07]. There is no difference in slopes between both cases. *p*_*s*_ denotes the p-value from Kruskal-Wallis test for slope difference. [mean ± SEM in radians per unit position: CA1, -4.62 ± 0.08 vs -4.54 ± 0.10, *H*_(1)_ = 0.84, *p* = 0.3584; CA2, -4.48 ± 0.10 vs -4.57 ± 0.13, *H*_(1)_ = 0.06, *p* = 0.8117; CA3, -4.44 ± 0.12 vs -4.20 ± 0.14, *H*_(1)_ = 1.69, *p* = 0.1940]. (C) Histograms of spike phases from precession samples show higher spike phase for passes against *θ*_rate_. P-values are derived from Watson-Williams test for the difference between the circular means (shown as vertical bars) between the two cases [along-*θ*_rate_ vs against-*θ*_rate_, mean ± SEM in radians: CA1, 3.66 ± 0.04 vs 3.77 ± 0.05, *F*_(1,2073)_ = 2.93, *p* = 0.0871; CA2, 2.50 ± 0.02 vs 2.50 ± 0.02, *F*_(1,19271)_ *<* 0.01, *p* = 0.9867; CA3, 1.61 ± 0.02 vs 2.22 ± 0.03, *F*_(1,17151)_ = 285.63, *p* = 1.5*e* − 63].

**Figure 4:**
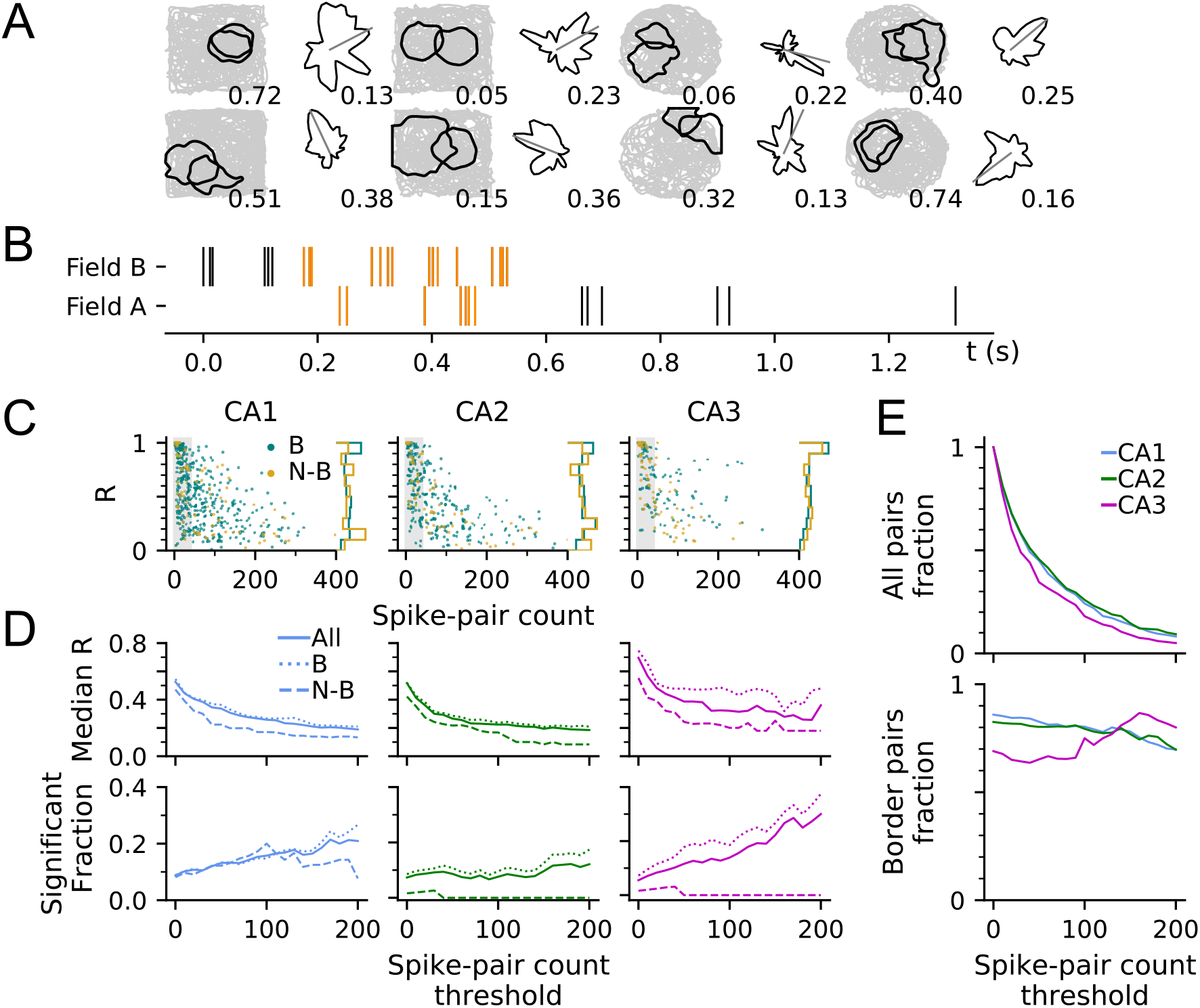
Pair correlations are highly directional in CA3. (A) Examples of pairs of place fields (black contour lines) and the directional tuning curves of paired spikes. Number below the contour plot indicate the amount of field overlap (calculated as 1 − *D*_*ks*_, where *D*_*ks*_ is 2D Kolmogorov–Smirnov distance between two place fields). Resultant vector length (*R*) of the directional distribution is printed below the tuning curve. (B) Raster plots of spike times in a pair of overlapping fields during traversal from field B to A. Admitted paired spikes with time difference *<* 0.08*s* between field A’s and B’s are colored in orange. (C) *R* and spike-pair count for the whole populations of border and non-border pairs in CA1, CA2 and CA3. Pairs with spike-pair counts below 40 are excluded in our statistical comparisons, indicated by the shaded region. (D) Median *R* (top) and fraction of significantly directional pairs (bottom) by different spike-pair number thresholds. Spike pairs in all CA regions were significantly directional [CA1: 31*/*258 = 12.02% (Binomial test, *p* = 6.9*e* − 06), CA2: 17*/*181 = 9.39% (*p* = 0.0097), CA3: 9*/*88 = 10.23% (*p* = 0.0319)]. Comparing regions in terms of *R*, we found that CA3 has higher directional selectivity than CA1 [Kruskal-Wallis test: CA1 (*n* = 258) vs CA2 (*n* = 181) vs CA3 (*n* = 88), *H*_(2)_ = 10.82, *p* = 0.0045; *post hoc* Dunn’s test with Benjamini-Hochberg correction: CA1 vs CA2, *p* = 0.1413, CA2 vs CA3, *p* = 0.0030, CA1 vs CA3, *p* = 0.0315]. In CA1 and, different from the single spike results in Figure 1, also in CA3 directionality is induced by the proximity to the border in terms of resultant vector lengths [Kruskal-Wallis test: CA1, border (*n* = 217) vs nonborder (*n* = 41), *H*_(1)_ = 4.39, *p* = 0.0361; CA3, border (*n* = 56) vs nonborder (*n* = 32), *H*_(1)_ = 9.64, *p* = 0.0019]. (E) Fraction of all place fields (top) and border fields (bottom) by spike-pair count thresholds.

For illustration, we plotted the typical characteristics of phase precession for cases when the animal runs along (|*d*| *<* 30°) or against (|*d*| *>* 150°) the preferred rate direction (*θ*_*rate*_) by separately fitting a regression line for the phase-position relation to passes from all fields in these two cases (Figure 3B) and found that passes aligned to the opposite of best rate direction indeed have on average a significantly higher onset of precession than those with different directions in CA3 region, but not in CA1 and CA2. The average precession slopes do not differ between the two groups of passes.

The difference in onset phases between along-*θ*_rate_ and against-*θ*_rate_ passes is also corroborated by the phase histograms (Figure 3C), where in CA1 and CA3 the against-*θ*_rate_ group exhibits a significant shift to later phase as compared to along-*θ*_rate_ group.

Thus, at least in CA3, phase precession tends to start from a higher phase when the rat’s running direction aligns with the opposite of preferred rate direction corroborating that phase precession exhibits directional modulations.

### 3.4 Directional selectivity in paired place fields

Phase precession is often considered a single-cell reflection of place cell sequences during theta (Dragoi and Buzsaki, 2006; Foster and Wilson, 2007; Feng et al., 2015; Leibold, 2020), and as such it should show up in peak lags of pair correlation functions, too (Dragoi and Buzsaki, 2006; Huxter et al., 2008; Geisler et al., 2010; Schlesiger et al., 2015). In 2-dimensional environments, such correlation lags have been shown to flip signs depending on the order in which a trajectory samples the place fields (Huxter et al., 2008), arguing for strong external (behavioral/sensory) drive of sequence structure. We therefore hypothesized that if phase precession reflects sequence firing, correlation lags should also be tuned to certain directions. To test our assertion, we first identified overlapping pairs of place fields (see examples in Figure 4A) and included only spike pairs that are spaced at most 80 ms in time (Figure 4B). This criterion allowed us to admit only the spike pairs that form part of a putative theta firing sequence.

Overall, directionality results are very comparable between spike pair and single spike analysis from Figure 1: We observe significant directionality in all subregions and a higher median *R* in CA3 (Figure 4D; see caption for statistics). Most importantly, however, directional tuning of pairs is much higher (in terms of median *R*) than of single spikes (Kruskal-Wallis test for median *R* difference between single spikes and spike pairs at spike count threshold 40: CA1, single (*n* = 800) vs pair (*n* = 258), *H*_(1)_ = 153.84, *p* = 2.5*e*−35; CA2, single (*n* = 521) vs pair (*n* = 181), *H*_(1)_ = 113.50, *p* = 1.7*e* − 26; CA3, single (*n* = 396) vs pair (*n* = 88), *H*_(1)_ = 78.70, *p* = 7.2*e* − 19), and thus we conclude that spike correlations in theta sequences induce additional directionality in line with our initial hypothesis.

Again, separating field pairs into border and non-border, we find that pair directionality in CA3 is also border sensitive (see caption for statistics) in contrary to single spike directionality. Particularly, border-sensitive CA3 pairs extend to high *R* values even for spike-pair counts *>* 100 (see Figure 4C). These findings suggests that border sensitivity in CA3 is specifically tied to the correlation structure, whereas in CA1 it’s mostly inherited from the directional firing rates.

### 3.5 Directionality of pair correlation

Since the pair firing rate showed region-dependent differences, we hypothesized that these differences should also transfer to spike timing correlations (Figure 5A). Huxter et al. (2008) already reported that spike correlation lags in CA1 depend on path direction and thus show a strong extrinsic (sensory / behavioral) dependence and so we followed their approach and confirmed their main results for all three CA regions: Correlation lags decrease as the field pairs’ overlaps increase and the sign of the lags flip if the path direction is reversed (Figure 5B).

**Figure 5:**
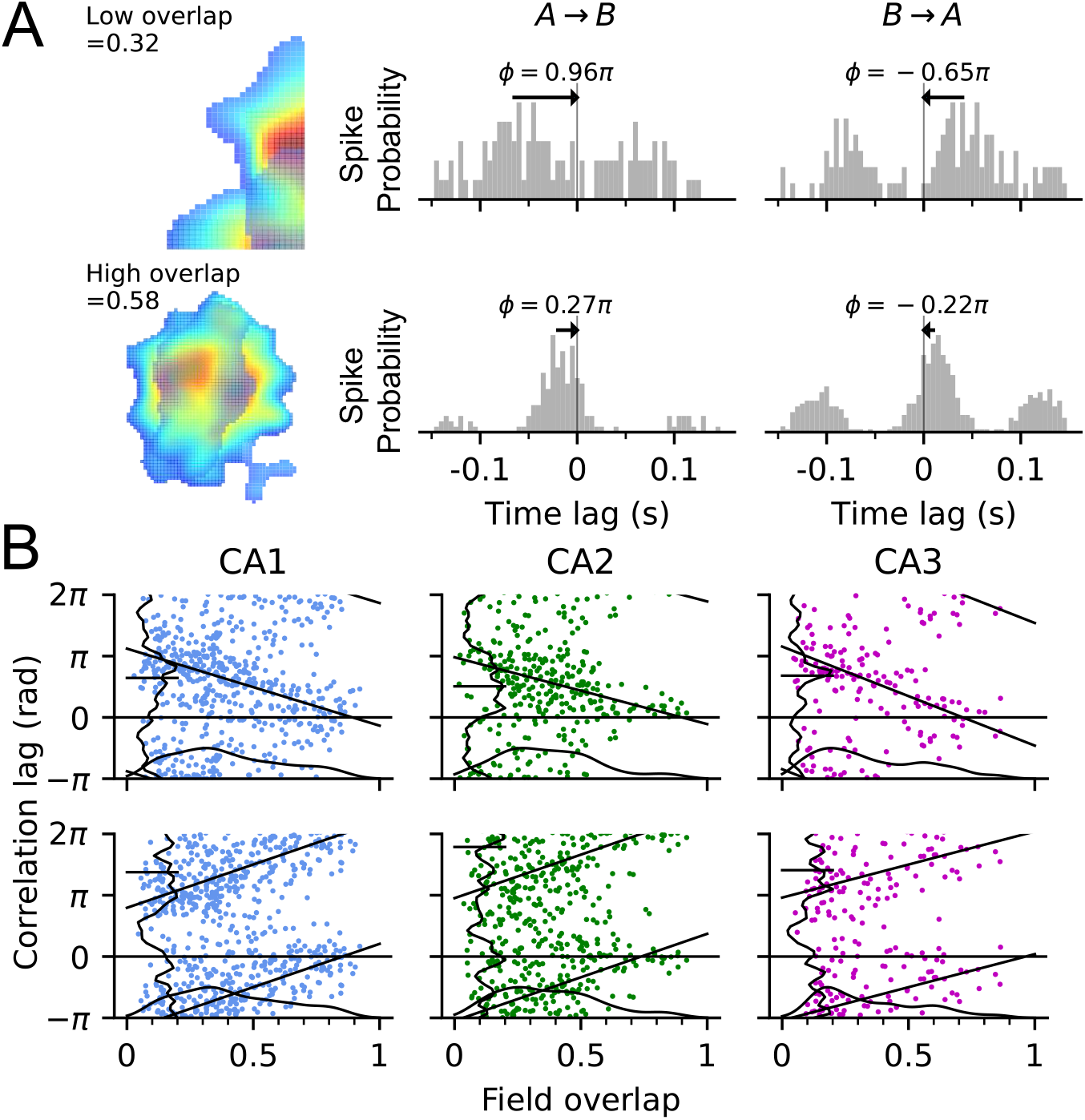
Strong extrinsic effect on correlation lag in all CA regions. (A) Top row: Examples of a field pair with low overlap (left) and its cross-correlograms when the animal runs from field A to field B (middle) and in the opposite direction, from B to A (right). The spike correlation lag *f*, indicated by the distance of the closest peak to mid-line, generally flips sign when the direction is reversed. Bottom row: Example of a field pair with high overlap. Note that the magnitude of correlation lag is smaller than the high-overlap example. (B) The higher the field overlap, the larger the spike correlation lag. Relation between spike correlation lags and the amount of field overlap for all pairs in CA1, CA2 and CA3 when the animal runs from field A to B (Top) and from field B to A (bottom). Black straight line shows linear-circular regression line [Direction A to B: CA1, *r*_(412)_ = −0.43, *p <* 1.0*e* − 63; CA2, *r*_(260)_ = −0.39, *p* = 2.9*e* − 09; CA3, *r*_(134)_ = −0.48, *p* = 6.6*e* − 08. Direction B to A: CA1, *r*_(414)_ = 0.49, *p <* 1.0*e* − 63; CA2, *r*_(280)_ = 0.17, *p* = 0.0030; CA3, *r*_(121)_ = 0.39, *p* = 1.3*e*−05]. Vertical and horizontal curves are the marginal distribution of spike correlation lags and field overlap respectively.

However, a closer inspection of the correlation functions reveals that they often do not have a clear single peak and thus we devised a new approach taking into account the symmetries of the full correlation function. Field pairs whose activities rely on intrinsic dynamics and are insensitive to sensory stimulus should show a similar shape of the correlation function regardless of reversing the path direction. In contrast, the correlation lags of extrinsic pairs should flip sign as an effect of direction reversal (for single pass example, see Figure 6A, C). Based on this principle, we were able to quantify “intrinsicity” and “extrinsicity” of a pair, using the overlap of the correlation functions of both pass directions (original and flipped: extrinsicity; original and original: intrinsicity; Figure 6B and D, also see Materials and Methods). Pairs with higher extrinsicity than intrinsicity are classified as extrinsic, and vice versa.

**Figure 6:**
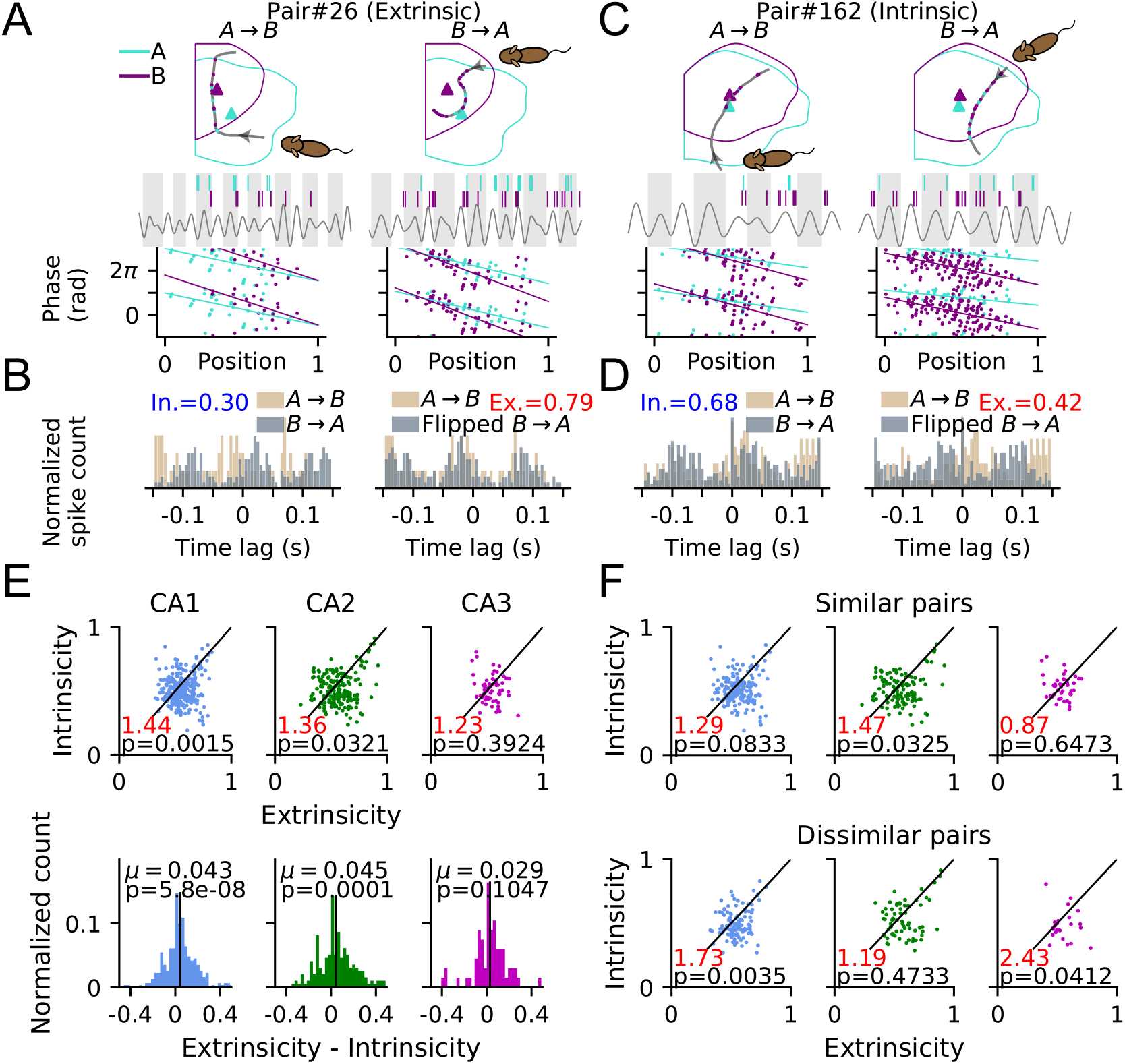
Place field correlations reveal region-specific extrinsic and intrinsic contributions. (A) Top: Illustration of theta sequence from an example extrinsic pair during a single pass when the rat runs from field A to B (*A* → *B*, left). Place cell A (sky blue) fires ahead of B (purple) in each theta cycle (gray and white shaded intervals). When the trajectory direction is reversed (*B* → *A*, right), place cell B fires ahead of A. Bottom: Phase-position relations of spikes for the example pair in (A), pooling from all passes in condition *A* → *B* (left) and *B* → *A* (right). (B) Intrinsicity (blue text) is computed by Pearson’s correlation (see Materials and Methods) between the cross-correlograms of two running directions (*A* → *B* and *B* → *A*, left panel), while the computation of extrinsicity (red text) takes the cross-correlogram of *A* → *B* and flipped cross-correlogram of *B* → *A* (right panel). (C) Same as (A), but an intrinsic pair. Cell B fires ahead of A even in the direction *A* → *B*. (D) same as (B), but with higher intrinsicity than extrinsicity. (E) Top: Scatter plots of intrinsicity versus extrinsicity for all field pairs. The diagonal line represents the decision boundary: pairs above it are classified as intrinsic pairs, and pairs below it as extrinsic pairs. Ratio of extrinsic pairs to intrinsic pairs (red number) and p-values (one-way Chi-square test) indicate significant bias towards extrinsicity for CA1. Bottom: Histograms of differences between extrinsicity and intrinsicity. Mean *µ* (black bar) and p-value of the t-test of mean versus 0 suggest a bias towards exrinsicity for all brain regions. (F) Intrinsicity versus extrinsicity for pairs with similar (difference *<* 90°, top row) and dissimilar (*>* 90°, bottom row) best rate angles. Detailed statistics are reported in the text.

The ratios of extrinsic to intrinsic field pairs in CA1 and CA2 are significantly different from the expected equality, while CA3’s is not (Figure 6E top, one-way Chi-square test for equal proportion of extrinsic and intrinsic pairs: CA1, 184 : 128 = 1.44, 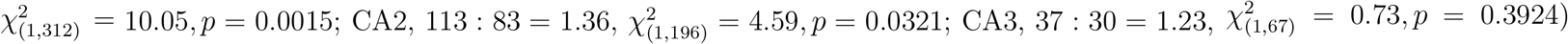. As a further measure for a bias towards extrinsicity or intrinsicity that also takes into account the amount of ex(in)trinsicity, we computed their difference, and found that all regions exhibit a significant bias towards extrinsicity except in CA3 (Figure 6E bottom, Student’s t-test for extrinsicity-intrinsicity with expected value of 0: CA1, mean=0.0434, *t*_(311)_ = 5.56, *p* = 5.8*e* − 08; CA2, mean=0.0453, *t*_(195)_ = 3.92, *p* = 0.0001; CA3, mean=0.0291, *t*_(66)_ = 1.64, *p* = 0.1047). We therefore conclude that the pair correlations in CA1 and CA2 demonstrate a strong dependence on path directionality but not significantly so in CA3.

The most parsimonious explanation for intrinsic pair correlation structure would be to assume that place cells are bound into rigid sequences that play out independent of running direction. In such a scenario place field firing should be highly directional, and pairs with similar preferred direction should reveal more of the intrinsic correlation structure than pairs with opposite preferred direction. We thus further separated pairs into those with similar (angle difference *<* 90°) and dissimilar (*>* 90°) best rate angles (Figure 6F) and found that, pooling over all CA regions, the similar pairs indeed have lower extrinsic-intrinsic ratio than dissimilar pairs (similar=203:158=1.28, dissimilar=131:83=1.58, Fisher’s exact test: *p* = 0.0353). Resolving for the different CA regions, the effect was significant in CA1 (similar=108:84=1.29, dissimilar=76:44=1.73, *p* = 0.0441) and CA3 (similar=20:23=0.87, dissimilar=17:7=2.43, *p* = 0.0333) but not in CA2 (similar=75:51=1.47, dissimilar=38:32=1.19, *p* = 0.0930). Thus place field pairs with similar directional tuning may contribute more to the activation of intrinsic sequence activation at least in CA1 and CA3.

Contrary to the extrinsic pairs, the intrinsic pairs are invariant to sensory inputs and maintain their firing order even when they are reversely sampled (see Figure 6C for single pass example). A possible mechanism could be that intrinsic pairs are asymmetrically connected and bias the generation of theta sequences in one direction. We thus hypothesized that the pair correlation structure reflects two contributions, extrinsic sequences which are produced by sensory inputs and flip order with the running directions, and, in addition, intrinsic sequences produced by the internal connections in a fixed order.

In order to dissociate the intrinsic sequences from the extrinsic ones, we focused on intrinsic pairs and for those separated the running trajectories into two groups: either they align with (“Same” group) or are opposite to (“Opposite” group) the intrinsic directional bias that is identified via their cross-correlation signals: A directional bias of *A* → *B* would mean the correlation signal always has a peak at negative time lag even when the animal traverses from B to A (Figure 7A). Similarly, a directional bias of *B* → *A* is determined by having both correlation peaks at positive time lag (Figure 7B). In the “Same” passes, both extrinsic and intrinsic sequences could be present and hardly distinguishable, since they have the same firing order. While in the “Opposite” passes, the occurrence of extrinsic sequences is less, as indicated by the lack of reversal of the cross-correlation signal, and hence, the theta sequences would be more representative of the underlying intrinsic firing order. Therefore, a comparison between “Same” and “Opposite” passes could reveal the difference between extrinsic and intrinsic sequences. More specifically, we asked whether phase precession, as a potential single-cell reflection of a theta sequence, would differ in onsets, slopes and phase distributions between “Same” and “Opposite” running directions.

**Figure 7:**
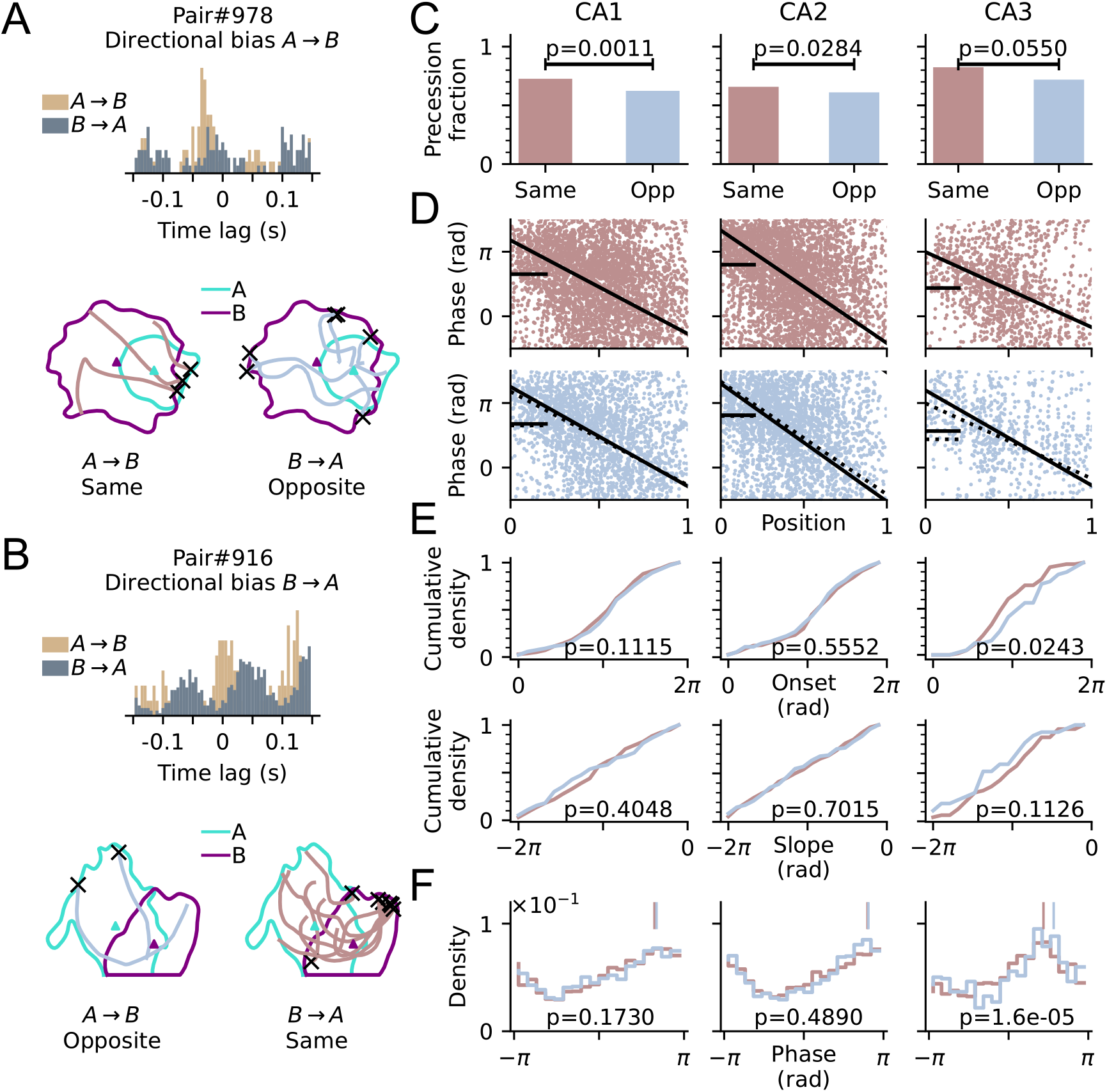
Prospective representation is revealed in CA3 intrinsic pairs (A) Top: Correlogram of an example pair with directional bias *A* → *B* (A always fires before B). Bottom: Passes that align with the directional bias *A* → *B* (left, brown, denoted as “Same”), and those are opposite to the directional bias (right, blue, denoted as “Opposite”). Origins of passes are marked as black crosses. (B) An example pair with directional bias *B* → *A*. (C) Fraction of passes that are precessing for “Same” and “Opposite” (abbreviated as “Opp”) conditions. P-values are derived from Fisher’s exact test. (D) Top: Phase-position relation for all precessing “Same” passes. Black solid curve marks the average of all individual linear-circular fits from each precession, and black bar marks the marginal mean phase. Bottom: Same as the top panel, but for “Opposite” condition instead. Linear-circular fit and marginal mean phase for “Same” passes from the top panel are also shown in dashed lines here for comparison. (E) Top: Cumulative density of onsets for all precessing “Same” and “Opposite” passes. Colors as in (C). P-values are derived from Watson-Williams test for differences in onset phases. Bottom: Cumulative density of slopes. P-values are derived from Kruskal-Wallis test for differences in slopes. (F) Distribution of spike phases for all precessing “Same” and “Opposite” passes. P-values are derived from Watson-Williams test for differences in mean phases (shown as vertical bars). See text for detailed statistics.

In line with our initial hypothesis, we found that precession is more likely to occur when passes align with the directional bias (Figure 7C). The “Same” condition has a higher fraction of precession than “Opposite” condition in all CA regions, but only reaching significance in CA1 and CA2 (Fisher’s exact test: CA1, *p* = 0.001; CA2, *p* = 0.0284; CA3, *p* = 0.0550). Considering only the precessing passes, there is no difference in precession slopes between “Same” and “Opposite” (Figure 7E bottom, Kruskal-Wallis test: CA1, Same (*n* = 236) vs Opp (*n* = 174), *H*_(1)_ = 0.69, *p* = 0.4048; CA2, Same (*n* = 213) vs Opp (*n* = 164), *H*_(1)_ = 0.15, *p* = 0.7015; CA3, Same (*n* = 61) vs Opp (*n* = 39), *H*_(1)_ = 2.52, *p* = 0.1126), but the phase onset of precession exhibits clear regional differences. CA3 has higher phase onset when the passes are opposite to the directional bias (Figure 7E top, Watson-Williams test: CA3, *F*_(1,98)_ = 5.24, *p* = 0.0243), while such difference is not significant in CA1 (*F*_(1,408)_ = 2.54, *p* = 0.1115) and CA2 (*F*_(1,375)_ = 0.35, *p* = 0.5552). Consistently, pooling the spikes from all precessing passes, the spike phase distribution from “Opposite” passes shows a significant shift to later phases as compared to “Same” passes only in CA3 (Figure 7F, Watson-Williams test: CA3, *F*_(1,2002)_ = 18.76, *p* = 1.6*e* − 05) but not in CA1 (*F*_(1,6017)_ = 1.86, *p* = 0.1730) and CA2 (*F*_(1,6272)_ = 0.48, *p* = 0.4890).

According to our hypothesis that extrinsic and intrinsic sequences are played out in parallel and that intrinsic pairs result from a suppression of extrinsic sequences in one direction, the association between intrinsic sequences and later spike phases, should be more strongly visible in intrinsic pairs than in extrinsic pairs. We thus also inspected the phase distributions in extrinsic pairs. Since, by definition, extrinsic pairs do not have a directional bias, we resorted to compare the spike phase distributions from all precessing passes between extrinsic and intrinsic pairs, regardless of their pass directions. The upward shift of spike phases in intrinsic sequences (if they include more intrinsic pairs), should then show up as a higher marginal spike phase than that of extrinsic pairs. Statistical analysis confirms this prediction and shows that precessing passes from intrinsic pairs indeed exhibit higher spike phases than extrinsic pairs in CA2 and CA3, but not in CA1 (Watson-Williams test, mean ± SEM in radians: CA1, Ex 2.07 ± 0.018 vs In 2.03 ± 0.026, *F*_(1,15816)_ = 0.82, *p* = 0.366; CA2, Ex 2.30 ± 0.020 vs In 2.59 ± 0.032, *F*_(1,9227)_ = 45.28, *p* = 1.8*e* − 11; CA3, Ex 1.20 ± 0.035 vs In 1.44 ± 0.044, *F*_(1,4232)_ = 13.68, *p* = 0.0002).

Our results support our hypothesis that intrinsic pairs display two types of sequences, extrinsic and intrinsic ones, in the “Same” direction and predominantly intrinsic sequences in the “Opposite” direction. Trajectories “Opposite” to the directional bias of spike pair lags display later spike phases and higher onsets.

It might seem counter-intuitive that, when the intrinsic pairs are traversed in the opposite direction, sequences play out in a backward order, and yet there is phase precession. We therefore propose a possible explanation for the coexistence of intrinsic sequence and phase precession that is in line with our analysis results, by assuming that intrinsic pairs consist of a leading cell and an enslaved cell. The enslaved cell only fires with a delay after the leading cell was active. Consequently, their correlation structure is fixed in both directions. As the leading cell undergoes phase precession, the enslaved cell fires after the precessing spikes of the leading cell, and hence, also precesses with a phase shift (see Figure 8A for schematic illustration).

**Figure 8:**
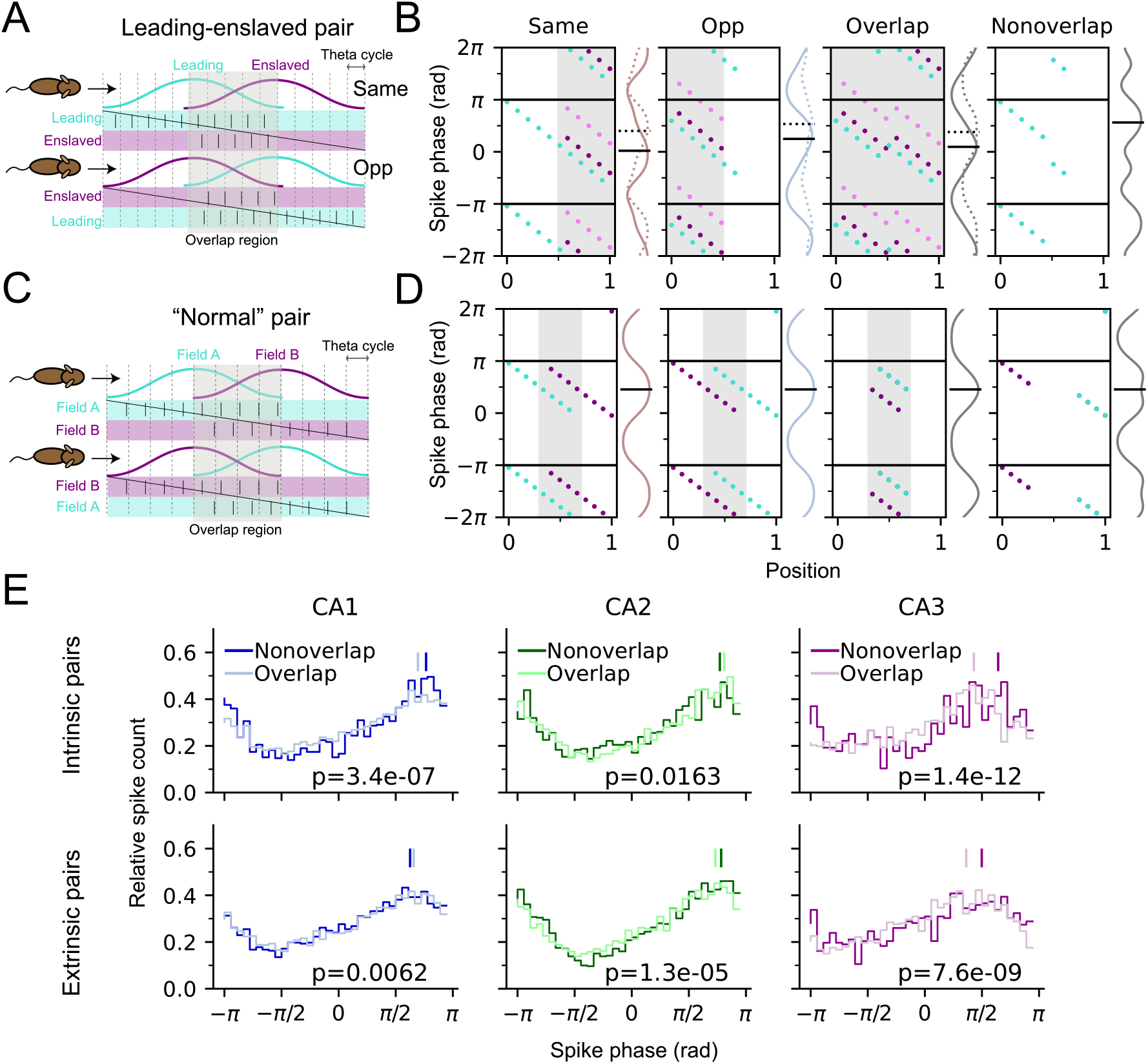
Co-occurrence of phase precession and intrinsic sequences. (A) Schematic illustration of how intrinsic sequences arise from enslaved spiking. The enslaved cells only fire at a fixed delay after the leading cell, causing a fixed temporal order even if the animal runs in the opposite direction (Opp, bottom row). This relation predicts that enslaved cells mostly fire in the overlap region of the two place fields (shaded area). Phase precession of leading (cyan dots) and enslaved (dark purple) cells plotted for Same, Opp, within and outside of overlap region, respectively from left to right. Data was generated artificially for illustration purposes. Marginal distribution of spike phase (solid curve) and its circular mean (solid black bar) are shown to the right of each panel. Light violet dots and dashed lines represent an alternative data set of enslaved spikes with larger phase shifts from the leading spikes. Note that the phase means of Opp passes and in non-overlap regions are higher than Same and overlap, respectively. (C) Schematics of theta sequences produced by “Normal” pairs without enslaved spikes. Note that the firing order is reversed when the pair is traversed in the opposite direction and both cells fire symmetrically in the non-overlap region. (D) Phase precession in “normal” pairs. Note that there is no difference in marginal phase means across all conditions. (E) Phase distribution of precession spikes in Overlap and Nonoverlap regions for intrinsic (top) and extrinsic (bottom) pairs in CA1, CA2 and CA3. P-values are derived from Watson-Williams test comparing the circular difference of phases between non-overlapping regions and overlaps. CA3 shows a larger effect in the phase difference as compared to CA1 and CA2. See text for detailed statistics.

The first prediction of the enslavement hypothesis is that the enslaved cell only fires within the overlap region between the two fields. As a result, the “Opposite” condition would have a higher spike phase than the “Same” condition, since both leading and enslaved spikes are present in the prospective cycles at the beginning of the field (see Figure 8B, left panels). While in “Normal” pairs, without enslaved spikes (Figure 8C), there would be no phase difference between “Same” and “Opposite” (Figure 8D, left panels). The prediction is compatible with our experimental findings in Figure 7, where the marginal phase mean and onset in the “Opposite” condition are higher than in the “Same” condition, indicating a possible contribution of enslaved spikes to the intrinsic structure of CA3.

Another assumption in the enslavement hypothesis illustrated in Figure 8 is that the precession of leading cells in the “Opposite” condition would start at a lower onset and last for a shorter number of cycles than in Same, due to the diminished forward recurrent connection when the pair is traversed in the opposite order. It would lead to the second prediction that the mean phase in the region where place fields are non-overlapping should be higher than within the field overlap (Fig. 8B, the right panels). On the contrary, the “Normal” pairs would show no phase difference between overlap and nonoverlap regions as phase precession is symmetrical in both directions (Figure 8D, right panels).

The prediction of higher overlap phase in nonoverlap regions from the enslavement hypothesis can be tested by comparing the phase distributions between overlap and nonoverlap regions in the experimental data (Figure 8E top row). We found that in CA3 intrinsic pairs, nonoverlap spiking indeed has a higher mean phase than in the overlap (mean ± SEM in radians: Nonoverlap 2.02 ± 0.07 vs Overlap 1.35 ± 0.04, Watson-Williams test, *F*_(1,2745)_ = 50.69, *p* = 1.4*e* − 12). A similar trend is also observed in CA1 intrinsic pairs, but the difference is smaller than in CA3 (Nonoverlap 2.41 ± 0.03 vs Overlap 2.18 ± 0.02, *F*_(1,9905)_ = 26.03, *p* = 3.4*e*−07). These results support that in CA1 and CA3, the intrinsic structure is generated by enslaved spikes. As a control, we also inspected the nonoverlap-overlap phase differences in the extrinsic populations (Figure 8E bottom row). In CA3, the phase difference has decreased as compared to its intrinsic counterpart (CA3 extrinsic: Nonoverlap 1.57 ± 0.05 vs Overlap 1.14 ± 0.04, *F*_(1,3813)_ = 33.53, *p* = 7.6*e* − 09), while in CA1, there is even a higher mean phase in the Overlap region (CA1 extrinsic: Nonoverlap 1.97 ± 0.03 vs Overlap 2.06 ± 0.02, *F*_(1,17523)_ = 7.48, *p* = 0.0062), indicating that enslaved spiking is less involved in the extrinsic pairs.

Intrinsic sequences and phase precession are thus not contradicting each other, if we assume that one field exhibits only enslaved spikes. The existence of enslaved spikes can explain both higher spike phase in the opposite traversal direction as well as the non-overlap area, which are both in agreement with the data.

#### Relation between pair correlation and phase precession

Following up the hypothesis from Figure 8 that intrinsic pairs are resulting, at least partly, from enslaved spikes, we suggested that both phase precession and pair correlations contribute different aspects to theta sequences, as already proposed previously (Middleton and McHugh, 2016). We thus finally asked whether we can identify a direct relation between the intrinsicity/extrinsicity property and the directionality of phase precession in the data, to further corroborate our enslavement hypothesis. To this end, we distinguished trajectories that are either parallel (*<* 90°) or opposite (*>* 90°) to the best rate angles of the fields of a pair with parallel best rate directions (see schematic illustration in Figure 9A) and included only passes with phase precession in both fields. Since the late spike phases appear to be associated with the passes opposite to the best rate angles in single fields (Figure 2), as well as the intrinsicity in CA3 (Figure 7), we hypothesized that the association should also transfer to field pairs. Our analyses show that, indeed, when the trajectories oppose both of the best rate angles, there is a higher proportion of intrinsic pairs contributing to phase precession (Figure 9B) and a higher marginal spike phase of co-occurring phase precession in CA3 (Figure 9C) than in the case when the trajectory runs in parallel to both best rate angles. These directional differences are not observed in CA1 pairs. CA2, surprisingly, shows the opposite trend that higher spike phases are more associated with the best rate direction, but there is no difference between the contributions of extrinsic and intrinsic pairs. The findings on CA3 corroborate that both correlation structure and phase precession are direction dependent in CA3, where the intrinsic field pairs seem to lack strong extrinsic drive in one running direction and thus would exhibit enslaved spikes at later spike phases when the running direction opposes their best rate direction. In extrinsic pairs the extrinsic drive seems strong in all running directions to overrule the intrinsic structure and allows cells to fire at earlier phases.

**Figure 9:**
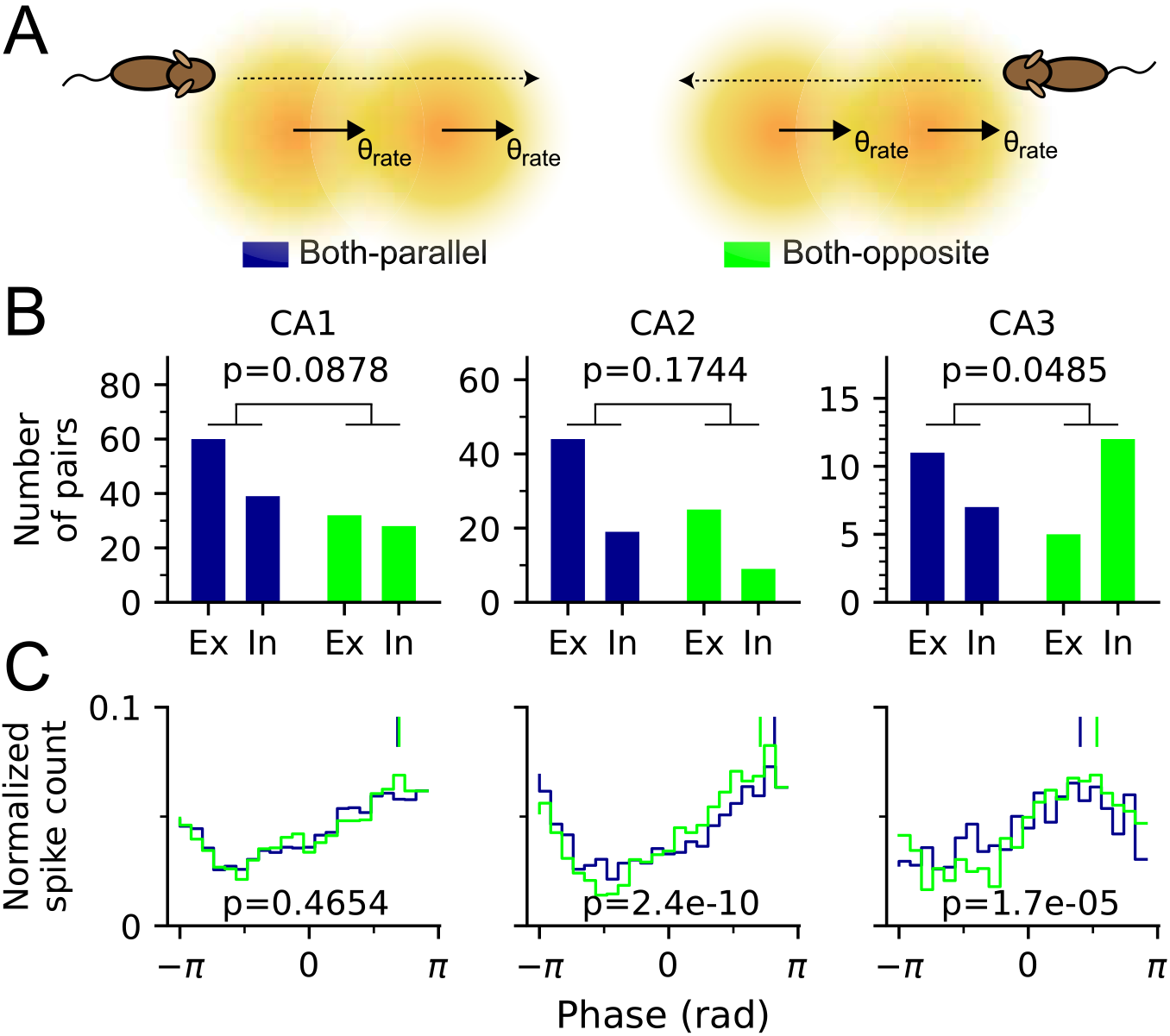
Field pairs in CA3 exhibit higher intrinsicity and fire at later spike phases in the direction opposite to the best rate angles. (A) Schematic illustration of how trajectories are divided into two groups, Both-parallel (left), if trajectories across a field pair deviate less than 90° from their best rate angles, and Both-opposite (right), if trajectories deviate more than 90° from their best rate angles. (B) Numbers of extrinsic and intrinsic pairs exhibiting phase precession in Both-parallel and Both-opposite cases. In CA3, there is significantly higher contribution from intrinsic pairs in Both-opposite case than in Both-parallel case. P-values are derived from Fisher’s exact test. (C) Phase distributions of phase precessing passes. The Both-opposite case in CA3 shows a significantly later spike phase than the Both-parallel case. Vertical bars mark the circular means. P-values are derived from Watson-Williams test [Both-parallel vs Both-opposite, mean ± SEM in radians: CA1, 2.15 ± 0.0252 vs 2.18 ± 0.0369, *F*_(1,7917)_ = 0.53, *p* = 0.4654; CA2, 2.57 ± 0.0321 vs 2.22 ± 0.0369, *F*_(1,4638)_ = 40.31, *p* = 2.4*e* − 10; CA3, 1.26 ± 0.0561 vs 1.67 ± 0.0609, *F*_(1,1841)_ = 18.55, *p* = 1.7*e* − 05].

### 3.6 Computational model as control

To further corroborate that the observed directionality effects in place field activity are not just an artifact of running trajectories or data analysis, we applied our analysis to simulations of a spiking model that does not include directional information as input (Romani and Tsodyks, 2015) but can be fed with the animal trajectories of our experiments (see Materials and Methods for simulation details). In brief, the model neurons integrate place specific inputs, theta-periodic inputs and symmetric recurrent connections with short-term synaptic depression. The recurrent connections give rise to omnidirectional phase precession but, owing to their symmetry, do not impose preferred intrinsic sequential activity. Since also the place-specific inputs are not directionally modulated, we reasoned that any directionality and any intrinsicity we find in the model must be artificial. Indeed, as expected, directionality analyses on the single field firing rates showed that the fraction of significantly directional simulated place fields does not exceed chance level of 5% (61/1024=5.96%, p=0.0936, Binomial test. Also see Figure 10A). Similarly, by pooling over all fields again, we did not find a significant correlation of phase precession per spike with the distance to preferred rate direction (Figure 10B, Spearman’s correlation coefficient *r*_*s*(1386)_ = −0.27, *p* = 0.0710). A further inspection on the distribution of preferred precession angles on a single field basis did not reveal a significant *π* Shift with respect to the preferred rate angles (Figure 10D, V-test against *π*: for all passes: *V*_(383)_ = −26.21, *p* = 0.9709; Figure 10E for low-spike passes, *V*_(174)_ = −30.61, *p* = 0.9995). After again separating the passes into groups of against-*θ*_rate_ (*n* = 105) and along-*θ*_rate_ (*n* = 361), with *>* 150° and *<* 30° angular difference in radians from the best rate angles respectively (Figure 10F), we consistently found no significant difference of their average precession curves (Kruskal-Wallis test for difference in slopes: *H*_(1)_ = 0.09, *p* = 0.7591; Watson-Williams test for difference in onsets: *F*_(1,464)_ = 0.01, *p* = 0.9415) and there was also no significant difference in the means of spike phases between group against-*θ*_rate_ and along-*θ*_rate_ (Watson-Williams test: *F*_(1,7639)_ = 2.78, *p* = 0.0953, see Figure 10G). Thus, as expected by model design, properties of phase precession in the model generally do not show a significant dependence on the animal’s heading directions.

**Figure 10:**
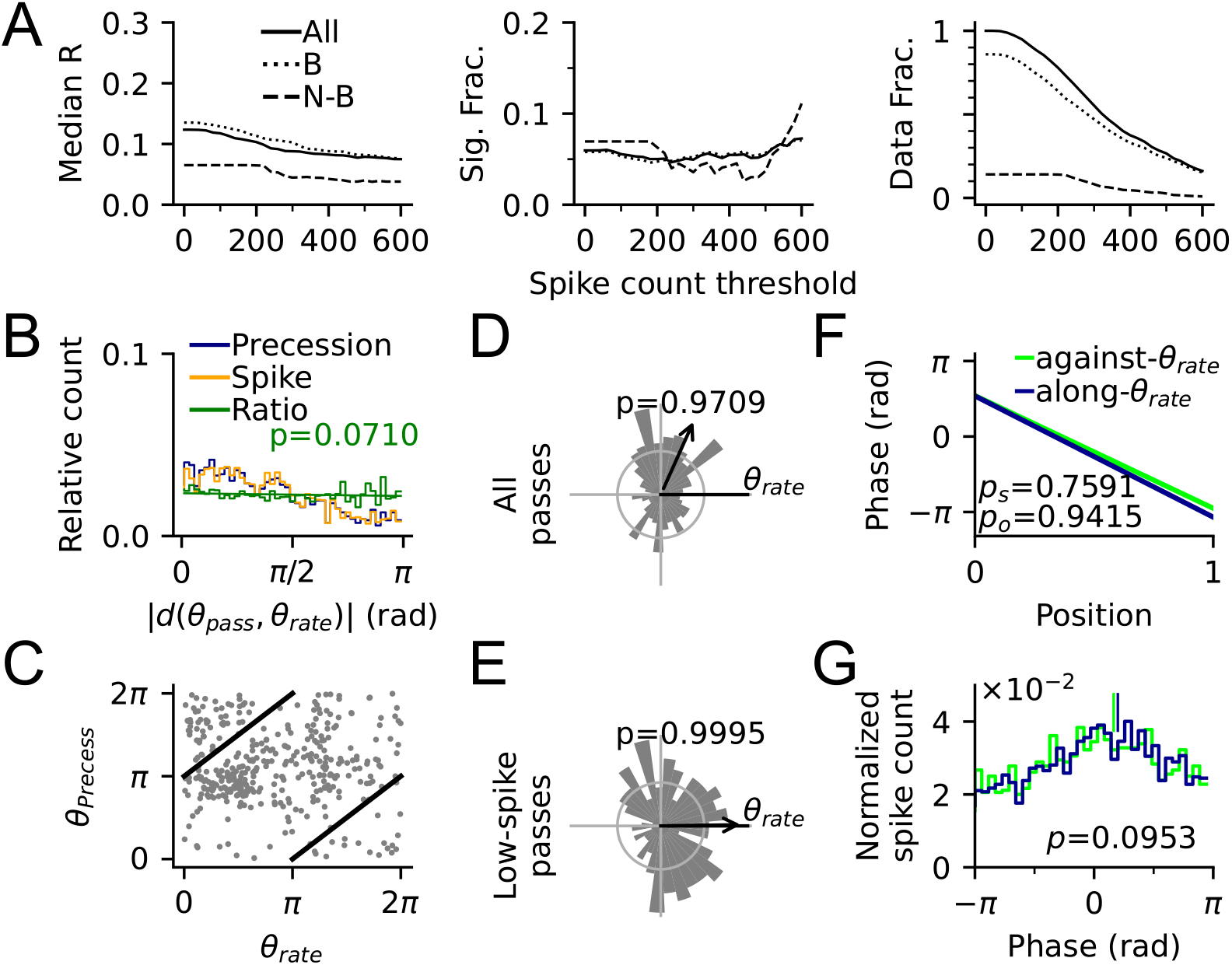
Simulation results using the model by Romani and Tsodyks (2015). (A) Directionality of simulated single fields. Median *R* (left) and fraction of significantly directional place fields (middle) by spike count thresholds for all (solid line), border (dotted) and non-border (dashed) fields. The rightmost panel shows the fraction of all, border and non-border place fields by spike count thresholds. (B) Distribution of precession incidences (dark blue) and spikes (orange) as a function of |*d*(*θ*_pass_, *θ*_rate_)|, the difference between pass direction and best rate direction of the place field. Ratio of blue and orange line in green shows no significant trend (Spearman’s correlation *r*_*s*(1386)_ = −0.27, *p* = 0.0710). (C) Scatter plot for the relation between best precession angle *θ*_precess_ and rate angle *θ*_rate_ of all place fields reveals no obvious structures. (D) Distribution of *θ*_precess_ directions of all precessing passes corrected to *θ*_rate_ shows no significant 180° difference of two best angles. (E) Distribution of *θ*_precess_ directions of low-spike passes. (F) Average precession slopes (phase-position curves) from precession samples against (green) and along *θ*_rate_ (blue). *p*_*s*_ and *p*_*o*_ are derived from Kruskal-Wallis test for slope difference and Watson-Williams test for onset difference respectively. (G) Distribution of spike phases for precession samples against and along *θ*_rate_. Vertical bars denote the circular means of the distributions. P-value is derived from Watson-Williams test comparing the difference of two circular means. Detailed statistics are reported in the text.

A virtue of the investigated Romani and Tsodyks (2015) model is that it contains recurrent synaptic connections and thus we asked to what extent they might explain intrinsic correlation structure observed in the data. We thus repeated directionality analysis at the level of field pairs on the simulated data. Similar to the single field results, the fraction of significantly directional pairs never exceed chance level in the model (169/3197=5.29%, p=0.2394, Binomial test, also see figure 11A), however, the dependence of correlation lags on place field distance can be well reproduced (Figure 11B), demonstrating that phase precession induces direction-dependent sequential firing of model place fields.

**Figure 11:**
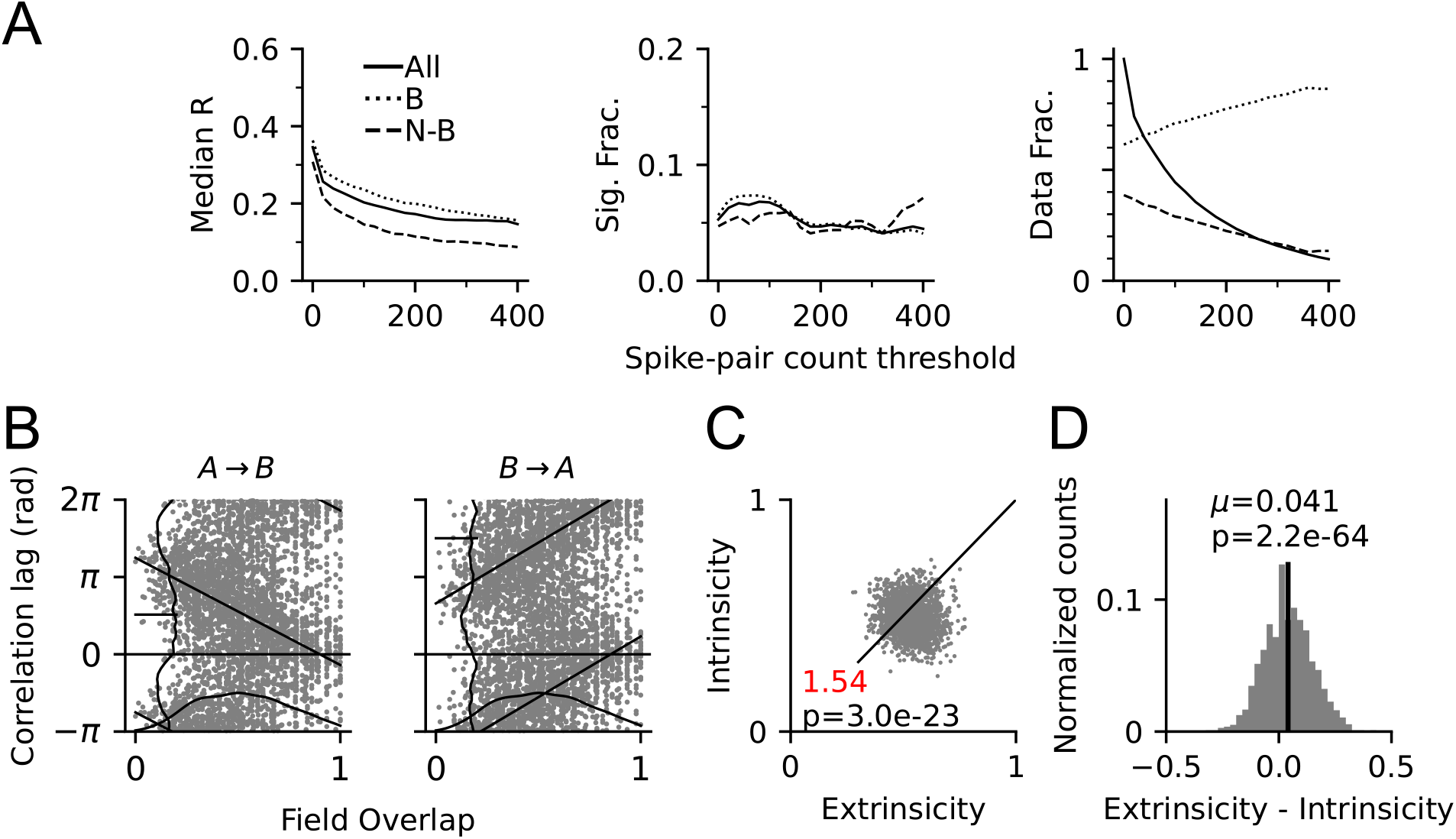
Directionality of pair correlations in model simulations. (A) Median *R* (left) and fraction of significantly directional pairs (middle) by spike count thresholds for all (solid line), border (dotted) and non-border (dashed) pairs. Right: fraction of all, border and non-border field pairs. (B) Relation of spike correlation lags and field overlap of all pairs for direction *A* → *B* (left) and *B* → *A* (right). Linear-circular regression line is depicted in black (*A* → *B, r*_(2659)_ = −0.38, *p <* 1.0*e* − 90; *B* → *A, r*_(2663)_ = 0.35, *p <* 1.0*e*−90). Vertical and horizontal curves show the marginal distributions. (C) Intrinsicity versus extrinsicity for all pairs (One-way Chi-square test of equal extrinsic-intrinsic ratio) and (D) Density of extrinsicity - intrinsicity (mean: black bar; p-value from Student’s t-test of mean versus 0) show a significant trend towards extrinsicity.

We next computed extrinsicity and intrinsicity of the model correlations and found a fraction of extrinsic pairs similar to CA1 (Figure 11C, Ex:In=1313:851=1.54, One-way Chi-square test for equal proportion of extrinsic and intrinsic pairs: 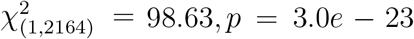) as well as the bias to extrinsicity (Figure 11D, Student’s t-test for extrinsicity-intrinsicity with expected value of 0: mean=0.0409, *t*_(2163)_ = 17.52, *p* = 2.2*e* − 64), suggesting that the balance between spatial inputs and recurrent connections in the Romani and Tsodyks (2015) model has an extrinsic bias most similar to CA1.

To conclude, although a network model with asymmetric weight couplings can reproduce the results of phase precession and correlation lags in a 2-dimensional environment including the extrinsic bias prominently observed in CA1 ensembles, further extension is still required to account for directional selectivity in place field networks and their effect on phase precession, specifically on the onset phase, and the less extrinsic dynamics of CA3.

## Discussion

We showed that theta-scale timing of place cell activity in all regions of the hippocampus proper exhibits directional modulations. This firstly applies to phase precession, which is most prominent (per spike) in the direction opposite to the one with the largest firing rate, and this effect is more prevalent and acute in CA3 than CA1 and CA2. In addition, best precession tuning in CA3 is associated with higher onset phases. Secondly, by inspecting directionality of spike pairs from two overlapping place fields, we showed that pair correlation exhibits higher directionality than single spikes from individual place fields. Furthermore, using cross-correlation analysis, we demonstrated that CA1 pair correlations are better explained by external sensory inputs induced by overt movement, than CA3 pairs. This suggests that CA3 place field correlations are more strongly intrinsically determined. In addition, a closer inspection into the intrinsic pairs revealed that the intrinsic sequences, which are invariant to the sampling order of place fields, are also associated with higher onset and spike phases in CA3 phase precession. Lastly, we employed the model by Romani and Tsodyks (2015) and showed that the interaction between directionality and theta activity, as described in the results above, cannot be solely explained by an omnidirectional generation mechanism of theta sequences. Nonetheless, the model shares a similar dependence on extrinsic information thus being more analogous to CA1 rather than CA3.

The directionality of place field firing in 2-d open environments has a long history of debate. While classical studies on the influence of landmark cues (Kubie and Muller, 1991), as well as early modelling work (Touretzky and Redish, 1996) clearly acknowledge the availability of directional information to the place cell system, there was disagreement about the degree to which this information becomes overt in place field activity (Muller et al., 1994; Anderson et al., 2006; Acharya et al., 2016). In the present study, we have found 10%-20% of significantly directional place fields, which is similar to previous reports (Markus et al., 1995; Acharya et al., 2016; Mankin et al., 2019), although directionality seems to strongly depend on the behavioral setting: Markus et al. (1995) found 80% of place cells with significant directionality in eight-arm mazes and 20% in an open circular platform. Acharya et al. (2016) also showed that the significant fraction of heading-direction modulated cells could vary as a function of the width of visual cue on the wall, since more place cells were head direction modulated near the border.

Most interestingly, and consistent with these previous reports, our analysis revealed that directionality of CA1 in familiar environments is heavily induced by proximity to the border, even at a high spike count threshold where the sampling bias is relatively small. However, this border effect is less prominent in simultaneously recorded CA3 place cells. One possible explanation is that there is no border-related directionality in novel environments but develops via learning processes, which is more difficult, and, hence slower in CA3, because of its recurrence. This idea has been suggested in a previous modelling study from Brunel and Trullier (1998) which demonstrated that place cells become less directionally selective through synaptic plasticity when they are visited in all directions (as in the non-border case), as compared to the border case when fields are visited only in a subset of directions. Also a second model by Navratilova et al. (2012) suggested that the firing rate of place cells is initially independent of directions, but develops to be directionally selective through experience and synaptic plasticity. Assuming that the same learning mechanism further differentiates between border and non-border directionality, the border difference would thus develop slower in CA3 due to its larger degree of recurrent connectivity. Both models thus support the idea that the differential directionality between border and non-border cells in familiar environments could be explained via plasticity induced by early experiences in novel environments. Our finding, however, that borders induces strong directionality of CA3 spike pairs (Figure 4), questions these models’ assumption that directionality is learned, but rather supports the idea that CA3 pair directionality reflects the directional imbalance of extrinsic and intrinsic activity (Figure 8).

The main conjecture of the present work was that inherent network dynamics could induce directionality of place cell spike timing. Through multiple lines of analysis, we repeatedly found that, in our data set, CA3 exhibits more hints to intrinsic dynamics than CA1. This was, e.g., suggested by their theta-scale correlations being more invariant to the trajectory. These findings are very much in accordance with anatomy: Pyramidal cells in CA3 project extensive recurrent collaterals in rodents and primates (Anderson et al., 2006; Amaral et al., 1984), while such recurrence is not so much obvious in the CA1 region (but see (Deuchars and Thomson, 1996) for rodents). However, a further possibility to explain CA3 bias towards intrinsic dynamics, could be its unique position in the mammalian hippocampal-entorhinal circuitry: Sensory information from entorhinal cortex layer II is projected to CA3 not only via the direct perforant pathway, but also via mossy fiber pathway of dentate gyrus (DG), contributing to an extra layer of internal processing through the granule and mossy cell loop. Indeed CA3 activity in rats with DG lesions showed reduced prospective firing in an eight-arm maze (Sasaki et al., 2018), which is strikingly consistent with our observed direction dependence of theta onset phases. The anatomical consistency of the enthorinal-hippocampal formation across mammals, as well as consistent reports on hippocampal phase precession in several mammalian species including chiroptera (Eliav et al., 2018) and primates (Qasim et al., 2021), let us speculate that the bias towards stronger intrinsic sequences in CA3 is a common feature within this animal class as a whole and specifically also expected to be seen for primates and humans despite their less precise spatial firing patterns (Ekstrom et al., 2003).

The directional selectivity of phase precession provides wide-spread support to our initial hypothesis that directional information should also express itself in hippocampal theta sequences, yet the anatomical foundations of this directional modulation can only be speculated about, so far. One possible explanation would be to assume that it is the direct entorhinal inputs to CA3 that induce the directional dependence of firing rate, which would be consistent with directional drive from postsubiculum and entorhinal head direction and conjunctive cells (Sargolini et al., 2006). According to Figures 2D and 7, this input would be reduced in passes opposite to best rate directions. Consequently, intrinsic CA3 activity would then mostly be explained by the indirect DG mossy fiber pathway, which was shown to induce prosepctive out-of-field spiking (Sasaki et al., 2018), potentially indicating enslaved spikes. In the best rate direction, according to our hypothesis, strong direct entorhinal input should induce earlier theta spike phases in addition to the DG-induced theta sequences. Conversely, against the best rate direction, DG-induced theta sequences would still play out invariantly despite the weaker entorhinal directional drive. In such a scenario, we would predict that intrinsic pairs would dominate phase precession properties over those extrinsically induced. A direct test of these predictions is provided in Figure 9, although the outcomes are consistent with our predictions the conclusiveness is limited owing to small sample sizes particularly in CA3. Yet the above hypotheses could be tested even more directly by acute differential suppression of entorhinal and DG pathways, which we would predict to selectively alter phase precession in the different running directions.

Intrinsic hippocampal correlations are incompatible with the interpretation of the hippocampus as a pure spatial map, but rather imply the existence of prestructure (Dragoi and Tonegawa, 2011). The degree to which intrinsic activity is expressed in familiar environments is, however, relatively small and clearly visible only in CA3, thus arguing for a relatively small role of intrinsic structure in supporting spatial navigation in familiar environments. The situation in novel environments, however, might be different with a stronger effect of prestructure on organizing hippocampal representations. This is in line with the finding that theta sequences only develop with time (Feng et al., 2015), since initially some pairs may predominantly fire in the “wrong” intrinsic order. The idea that – contrary to the models of Brunel and Trullier (1998) and Navratilova et al. (2012) – directional invariance develops in an experience-dependent way on top of prestructured sequences has important implications for theories of spatial memory formation, since it would favor the notion of sensory integration into existing temporal structure; cf. recent discussions of this idea in (Buzsáki and Tingley, 2018; Leibold, 2020).

## Acknowledgments

This work was funded by the German Research Association (DFG) under grant LE2250/13-1 and the NIMH under grant R01MH119179. The authors are grateful to Geoffrey W Diehl for help with the data transfer.

